# Waveform-based classification of dentate spikes

**DOI:** 10.1101/2023.10.24.563826

**Authors:** Rodrigo M.M. Santiago, Vítor Lopes-dos-Santos, Emily A. Aery Jones, Yadong Huang, David Dupret, Adriano B.L. Tort

## Abstract

Synchronous excitatory discharges from the entorhinal cortex (EC) to the dentate gyrus (DG) generate fast and prominent patterns in the hilar local field potential (LFP), called dentate spikes (DSs). As sharp-wave ripples in CA1, DSs are more likely to occur in quiet behavioral states, when memory consolidation is thought to take place. However, their functions in mnemonic processes are yet to be elucidated. The classification of DSs into types 1 or 2 is determined by their origin in the lateral or medial EC, as revealed by current source density (CSD) analysis, which requires recordings from linear probes with multiple electrodes spanning the DG layers. To allow the investigation of the functional role of each DS type in recordings obtained from single electrodes and tetrodes, which are abundant in the field, we developed an unsupervised method using Gaussian mixture models to classify such events based on their waveforms. Our classification approach achieved high accuracies (> 80%) when validated in 8 mice with DG laminar profiles. The average CSDs, waveforms, rates, and widths of the DS types obtained through our method closely resembled those derived from the CSD-based classification. As an example of application, we used the technique to analyze single-electrode LFPs from apolipoprotein (apo) E3 and apoE4 knock-in mice. We observed that the latter group, which is a model for Alzheimer’s disease, exhibited wider DSs of both types from a young age, with a larger effect size for DS type 2, likely reflecting early pathophysiological alterations in the EC-DG network, such as hyperactivity. In addition to the applicability of the method in expanding the study of DS types, our results show that their waveforms carry information about their origins, suggesting different underlying network dynamics and roles in memory processing.

**Author summary:** The entorhinal cortex and the dentate gyrus are regions of the brain’s hippocampal formation that play a crucial role in learning and memory. Their synchronous activation generates fast and high-amplitude electrophysiological patterns in the dentate gyrus, called dentate spikes (DSs). However, the functional role of DSs is still poorly understood. A technical limitation for the study of DSs is that their classification into types 1 and 2 is only possible through laminar profiles obtained by multicontact linear probes. In the present work, we propose a method to classify the different types of DSs through their waveforms, thus expanding the investigation of their significance in mnemonic processes by making possible their study in single-site recordings.

## Introduction

The hippocampal formation is critical for memory encoding, consolidation and retrieval [1–5]. Among its regions, the entorhinal cortex (EC) integrates polymodal sensory information and serves as an interface between the neocortex and the hippocampus [6]. On the other hand, the dentate gyrus (DG) acts as the main entry point to the hippocampus and is thought to function as a pre-processing unit that contributes to pattern separation [7–9]. The EC-DG communication occurs via the perforant path [10] which contains excitatory projections from EC layer II to the outer two-thirds of the DG molecular layer [11,12].

During slow-wave sleep, awake immobility and consummatory behaviors, the hippocampal local field potential (LFP) exhibits large-amplitude irregular activity (LIA) [13,14]. In such periods, coordinated discharges from EC to DG generate positive sharp patterns in the hilar LFP, known as dentate spikes (DSs) [15], which coincide with interregional gamma coherence [16] and seem to suppress the activity of CA3 and CA1 [15,17–19]. Concurrently, the complex of CA1 LFP patterns, referred to as sharp-wave ripples (SWRs), also occurs during LIA and has been extensively associated with memory consolidation processes [20–23]. While DSs may play a role in memory consolidation after associative learning tasks [24], disrupting DS-related processing has been found to enhance performance on tasks requiring DG pattern separation [25]. Nonetheless, further research is needed to fully understand the mechanisms and implications of DSs in brain network dynamics and cognitive processes.

In terms of electrophysiological signatures, DSs are fast (<30ms) and prominent (>1mV) transients linked to the synchronous discharge of granule cells (GCs), mossy cells (MCs), and inhibitory neurons [15,19,26]. Since the DS emergence depends entirely on the entorhinal inputs to the DG [15,27], two types of DSs are identified according to their origin – whether they arise from the discharges from the lateral (LEC) or medial EC (MEC) [15]. DSs are thus optimally classified based on the current source density (CSD) profile produced along the DG layers at the DS peak instant. This method is here called CSDbC (CSD-based classification). While the CSD of DS type 1 (DS1) exhibits a sink in the outer molecular layer (OML), which receives projections from the LEC, the CSD of DS type 2 (DS2) exhibits a sink in the middle molecular layer (MML), which receives projections from the MEC [15].

The use of linear probes with equally spaced electrodes is crucial for achieving an optimal DS classification through a detailed laminar profile of the DG. In principle, this requirement limits the DS-type analysis from other recording setups, such as single wire and tetrode [28]. The latter, for instance, provides a higher number of cells [29] and could expand the investigation of the functional role of each DS type. In this work, we developed a method called WFbC (waveform-based classification), enabling the classification of DSs based on their waveforms, thus extending their analysis to a broader range of recording setups.

Considering the classification obtained by CSD as the ground truth, we validated WFbC by analyzing the DSs of 8 mice from 3 distinct datasets with linear probes implanted in the dorsal hippocampus. Beyond calculating the WFbC accuracy, we also compared DS occurrence and morphological metrics of the DS types classified by both methods. In the process, we developed a new approach to measure the DS width based on waveform dynamics that was consistent across the DG layers, therefore serving as an alternative to traditional amplitude-based metrics [15,27,30].

Using the WFbC method on a dataset unsuitable for CSDbC due to its recording setup, we compared DS waveforms between apolipoprotein-ε3 (ApoE3) control and high Alzheimer’s risk apolipoprotein-ε4 (ApoE4) knock-in mice. We found consistently wider DS waveforms in young ApoE4 mice, mainly for DS2, indicating early entorhinal-hippocampal pathophysiological alterations before noticeable Alzheimer’s-linked cognitive behavior.

Taken together, the results demonstrate the feasibility of our method in classifying DSs through a single recording site, showing that the DS waveforms carry essential information about their origins and indicating unique activation dynamics of the DG network during each event type.

## Materials and methods

### WFbC validation datasets

To validate the WFbC method, we analyzed electrophysiological recordings from linear probes implanted in the dorsoventral axis of the hippocampus of 8 mice distributed across 3 datasets: dataset A with 1 mouse (A); dataset B with 3 mice (B1-3); and dataset C with 4 mice (C1-4). The animals and recording sessions were selected if they had at least 500 DSs of both types with a mean peak amplitude greater than 1 mV. Table 1 shows further details on the animals and their implants.

**Table 1.** Mice and electrophysiological implants used in WFbC validation.

| Mouse | Strain | Sex | Age (mo) | Linear probe | Implant coordinates (from bregma) | Study reference |
| --- | --- | --- | --- | --- | --- | --- |
| A | C57BL/6J (WT) | Male | 6 | NeuroNexus A1x32-Poly2-10mm-50s-177 | AP: -1.9 mm<br>ML: -1.2 mm | Senzai and Buzsáki, 2017 [26] |
| B1 | C57BL/6J (WT) | Male | 5 | NeuroNexus A1x32-6mm-50-177-H32_21mm | AP: -2 mm<br>ML: 1.7 mm | Lopes-dos-Santos et al., 2018 [31] |
| B2 | C57BL/6J (WT) | Male | 5 | Cambridge NeuroTech ASSY-236 H3 Chronic 64-Molex | AP: -2 mm<br>ML: 1.7 mm | Lopes-dos-Santos et al., 2023 [32] |
| B3 | C57BL/6J (WT) | Male | 5 | Cambridge NeuroTech ASSY-236 H3 Chronic 64-Molex | AP: -2 mm<br>ML: 1.7 mm | Lopes-dos-Santos et al., 2023 [32]. |
| C1 | C57BL/6J (PV-Cre) | Female | 4 | NeuroNexus A1x32-6mm-50-703-CM32 | AP: -1.95 mm<br>ML: 1.5 mm | Aery Jones et al., 2021 [30] |
| C2 | C57BL/6J (PV-Cre) | Female | 4 | NeuroNexus A1x32-6mm-50-703-CM32 | AP: -1.95 mm<br>ML: 1.5 mm | Aery Jones et al., 2021 [30] |
| C3 | C57BL/6J (SST-Cre) | Male | 4 | NeuroNexus A1x32-6mm-50-703-CM32 | AP: -1.95 mm<br>ML: 1.5 mm | Aery Jones et al., 2021 [30] |
| C4 | C57BL/6J (SST-Cre) | Female | 4 | NeuroNexus A1x32-6mm-50-703-CM32 | AP: -1.95 mm<br>ML: 1.5 mm | Aery Jones et al., 2021 [30] |

AP, anteroposterior; ML, mediolateral; PV, parvalbumin; SST, somatostatin; WT, wild type.

Mouse A was implanted with a 32-site polytrode that had channels arranged in two columns [26]. We focused on analyzing the raw LFPs (0-625 Hz) from the “140611” home cage session [33], whose electrode array position covered both DG blades. To ensure the DS amplitude profile along the dorsoventral axis, we specifically examined one-column channels, spaced 50 µm apart and numbered from 1 to 16.

Dataset B comprises raw LFPs (0-625 Hz) from periods when the mice were confirmed to be immobile within a familiar sleep box. The animal positions were monitored using a high-resolution camera, and the spatial data was processed in real-time by the Positrack software.

Mouse B1 was implanted with a 32-site linear probe with 50 µm inter-electrode spacing; its data were previously analyzed in [31]. Mice B2 and B3 were implanted with a 64-site linear probe with an inter-electrode spacing of 20 µm. To expedite the analysis process, we only considered the 40 lowest channels. Data from mice B2 and B3 have recently been analyzed in conjunction with data from mouse B1 elsewhere [32].

Dataset C is composed of the 0-300 Hz filtered LFPs from two parvalbumin-expressing (PV)-Cre mice and two somatostatin-expressing (SST)-Cre mice within their home cages, as described in [30]. Here we analyzed the concatenated LFPs from the sessions with vehicle (1% dimethyl sulfoxide in 0.9% sterile saline) administration [30]. The implanted 32-site linear probes had a distance of 50 µm between electrodes. Specifically for mouse C1, we considered the 18 lowest electrodes, since sites 9 and 11 were faulty. We also discarded its last session due to many visible artifacts over an extended period.

To provide consistency between the datasets for validation and application of the WFbC methods, we downsampled all LFPs to 1 kHz.

### WFbC application dataset

The dataset used to apply the WFbC method has been previously analyzed [34] and is publicly available on the CRCNS data-sharing website as hc-26 [35]. It comprises filtered hippocampal LFPs (0-300 Hz) sampled at 1 kHz and obtained from female apoE3-KI and apoE4-KI mice at rest across three different age groups: young, from 5 to 8 months; adult, from 9 to 11 months; and old, from 12 to 18 months. For this study, we merged the screen and replication cohorts mentioned in [34]. The criteria for selecting animals and channels required each session to have a minimum of 500 DSs with a mean peak amplitude greater than 1 mV. Due to this and to some individuals not surviving until old age, the composition of animals varied across the age groups (young apoE3-KI, n = 10; young apoE4-KI, n = 13; adult apoE3-KI, n = 11; adult apoE4-KI, n = 13; old apoE3-KI, n = 15; old apoE4-KI, n = 14).

The animals were implanted with 32-channel silicon probes (Neuronexus A4×8-5mm-200-400-703-CM32) targeting the right dorsal hippocampus (−1.95 mm AP and 1.5 mm ML from bregma) and were recorded in their home cages for 5 daily sessions, each lasting 60 min [34]. The probes had four shanks spaced 400 µm apart with eight electrode sites in each. The electrodes were vertically spaced 200 µm apart, exceeding a minimum distance to distinguish the current sinks between the middle and outer molecular layers and, thus, making the DS classification impossible via CSD.

### DS detection

DS detection consisted of finding the peaks of the filtered LFP (1-200 Hz, fourth-order Butterworth) that exceeded a threshold and were distant from each other for at least 50 ms. First, as a facultative step to mitigate artifact detection, we subtracted from the target LFP a reference signal, taken as the LFP from a channel outside the DG, with minimal potential variation but containing noise common to the probe. The detection threshold was set as seven times the median absolute amplitude of the filtered LFP, and the smaller peaks were first eliminated until the temporal distance condition was met. Then, to account for distortions introduced by the filtering process, we implemented a peak offset correction by defining each DS peak as the maximum value of the raw LFP within a 20-ms window around the peak identified in the filtered LFP. This time window was chosen based on the average DS width; at any event, most offsets were < 2 ms, thus using larger or shorter windows provide similar results. Finally, in a further attempt to avoid artifacts, we excluded the “DSs” whose peak amplitude fell out of the Tukey’s fences of the peak amplitude distribution (see Fig 1G-J), defined as [Q_1_ − 1.5(Q_3_ − Q_1_), Q_3_ + 1.5(Q_3_ − Q_1_)], where Q_1_ and Q_3_ are the first and third quartiles respectively.

**Fig 1.**
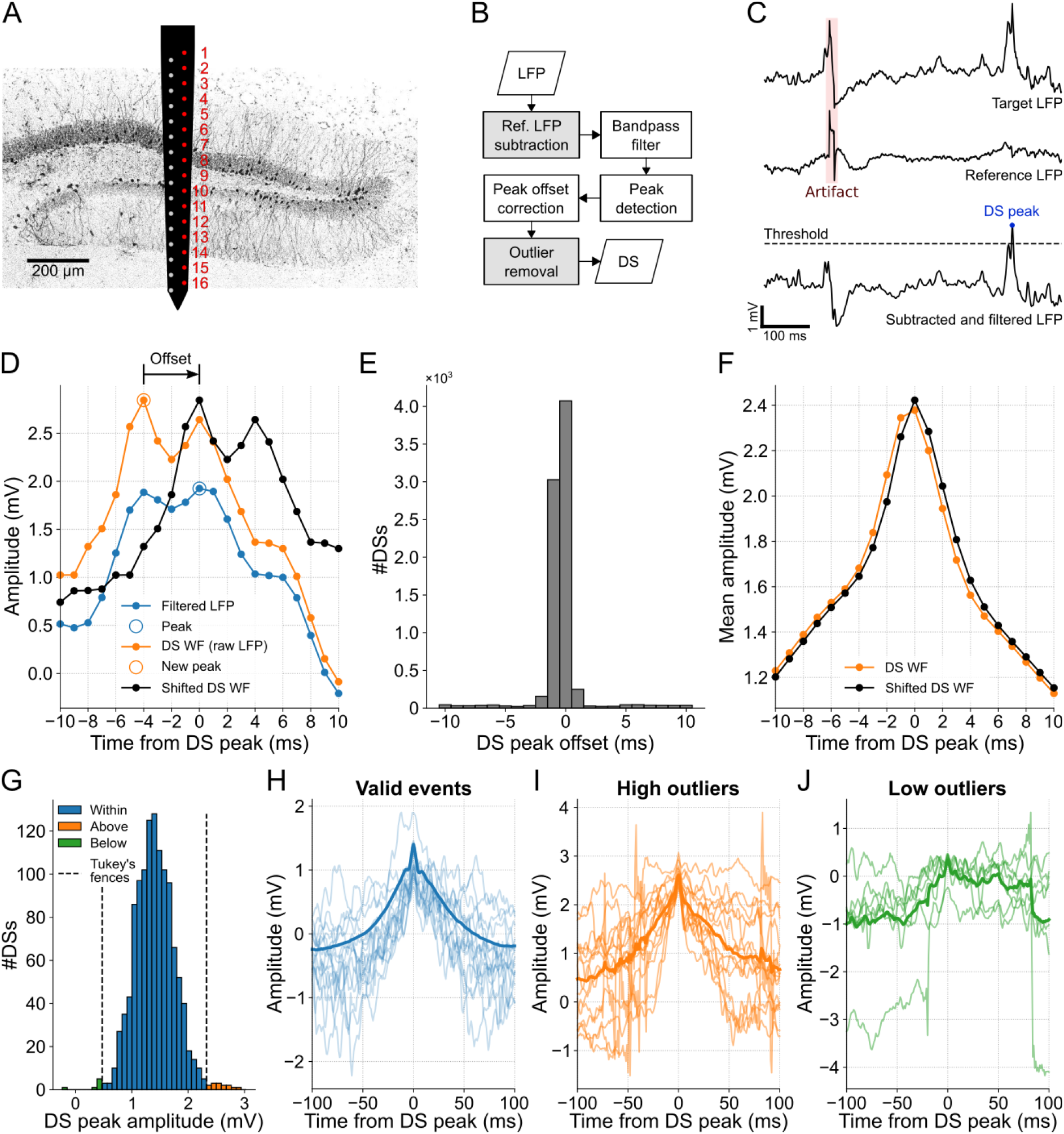
DS detection with artifact mitigation. **(A)** Schematics illustrating the two-column probe and its estimated position in mouse A. The electrodes used to obtain the DG laminar profile shown in subsequent figures are highlighted in red. **(B)** DS detection flowchart. Gray boxes represent artifact mitigation steps, which can be bypassed. **(C)** Example of DS peak detection. First, the reference LFP (middle trace) is subtracted from the target LFP (top trace) for artifact mitigation, a procedure followed by filtering (bottom trace). Finally, the peaks above the threshold are detected. **(D)** Example of peak offset correction. The DS waveforms are extracted from the raw LFP (orange trace) and centered around the peak detected in the filtered LFP (blue trace). If there is a higher peak in the raw signal within a 20-ms window, the signal is time-shifted to center the waveform on the new peak (black trace). **(E)** Offset distribution of the DS peaks detected in channel 9. **(F)** DS mean waveforms before and after the peak offset correction (orange and black traces, respectively). **(G)** DS peak amplitude distribution from channel 12 of mouse A. Peaks above (orange) and below (green) the Tukey’s fences (dashed lines) are ignored. **(H)** The first 15 waveforms (light blue traces) and the total mean waveform (blue trace) of the DSs whose peaks are within the Tukey’s fences in G. **(I,J)** Same as H, but for peaks above and below the Tukey’s fences. Note that the waveforms in I and J correspond to artifacts. DS: dentate spike; LFP: local field potential; WF: waveform.

### DS peak amplitude and scaled DS peak amplitude (metrics 1 and 2, M1 and M2)

DS waveform amplitude, measured from the raw LFP, varies along the DG lamina. For a proper comparison across animals, we calculated the z-score of the mean waveform from −200 to 200 ms relative to the peak, referred to as scaled mean waveform.

### DS rate (metric 3, M3)

The DS rate was estimated as the average rate computed over one-minute windows during strong delta (δ) oscillations. To infer delta activity, raw LFPs were subjected to the Empirical Mode Decomposition (EMD) method, which is used to analyze non-stationary signals [36]. Strong delta oscillations were then defined to occur when the power of the fifth intrinsic mode function (IMF-5) at the frequencies from 0 to 5 Hz corresponded to more than 90% of its full-band power (see Fig 2).

**Fig 2.**
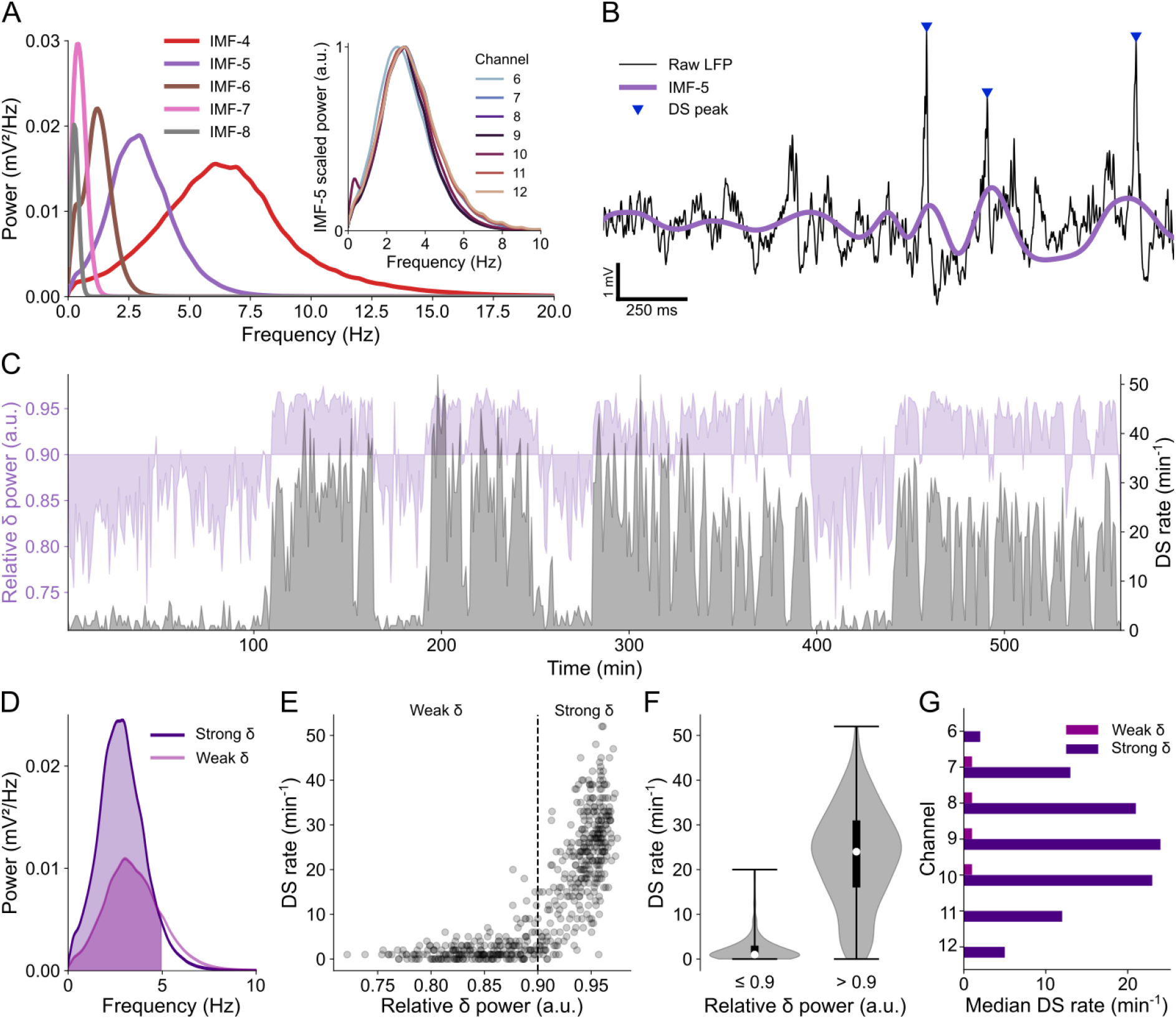
DS rate is correlated to the power of delta oscillations. **(A)** Power spectral densities (PSDs) of the main IMFs from the LFP of channel 9 in mouse A. The purple PSD (IMF-5) covers the delta frequency band. Inset: Normalized PSDs of IMF-5 across relevant channels. **(B)** Example of raw LFP from channel 9 (black trace) with DS peaks highlighted by blue triangles and the corresponding IMF-5 in purple. **(C)** Overlapping plots of relative delta power (in purple) and DS rate (in gray) in channel 9 over the entire recording session. **(D)** PSD of strong and weak delta waves. The colored fill indicates the delta range (0-5 Hz). **(E)** Scatter plot of the DS rate in function of the relative delta power. Each dot corresponds to the rate in a 1-minute bin and the dashed line separates weak and strong delta (relative power ≤ 0.9 and > 0.9 respectively). **(F)** DS rate distributions (violin plots) for weak and strong delta. Whiskers indicate the distribution range, white dots are the medians and thick black lines are the interquartile ranges. **(G)** Median DS rates during weak and strong delta in each relevant channel. DS: dentate spike; IMF: intrinsic mode function; LFP: local field potential.

### DS width and half-height width (metrics 4 and 5, M4 and M5)

The width was defined from the mean DS waveform. Particularly, we set the DS width as the temporal distance between the second upward concavities before and after the peak. The concavities were identified from the positive values of the second derivative of the mean waveform: the start width limit corresponded to the timestamp of the local maximum of the second derivative between −15 and −5 ms, while the end width limit was defined from the maximum between 5 and 15 ms. The width at half height was also calculated to verify the most stable metric among channels.

The LFP sampling period dictates the minimum non-zero difference between two width measurements, which is 1 ms for the sampling rate of 1 kHz. In order to get more accurate measurements, we reduced the sampling period by a quarter by resampling the mean waveforms to 4 kHz through a cubic spline interpolation before calculating the DS width.

### Classification of DSs based on their CSD profiles

The conventional method used to classify DSs is the CSDbC method, which relies on the analysis of their CSD profiles [15,32]. In this study, we used CSDbC as a reference to evaluate the performance of WFbC. Fig S1 shows the steps of the CSDbC method using the dentate spikes detected in channel 9 (hilus) of mouse A.

First, for the timestamp of each DS peak, we obtained a CSD profile across recording depths through the one-dimensional Poisson equation: CSD(y_i_) = −σ[V(y_i+1_) − 2V(y_i_) + V(y_i−1_)], where V is the electrical potential at the i^th^ electrode y, and the medium conductivity σ was assumed to be unitary for simplification. The CSD spatial profiles were concatenated, giving rise to a matrix of events by channels. After, we reduced the dimensionality of the CSD matrix using the first two principal components (PCs) (Fig S1B), which usually explained >85% of the variance. We then clustered the reduced CSD matrix into two groups using Gaussian Mixture Models (GMM) (Fig S1C; of note, we ended by utilizing only the first PC since it uniquely retained a bimodal projection of the data). Each event was assigned to the cluster with the highest probability. This means that all events in each cluster had probabilities > 0.5, with the majority > 0.9, demonstrating the robustness of the method. This consistently resulted in two distinct event classes with the most prominent sinks located in different regions of the molecular layer. The DS types were thus determined according to the position of the most prominent sink above the main source of the mean CSDs (Fig S1D). Specifically, the cluster with the main sink referring to the highest channel in the DG was classified as DS1, while the other cluster was classified as DS2. Finally, the possibility of having a single DS type was assessed by examining the second derivatives of the mean DS waveforms, as explained further below.

### Classification of DSs based on their waveforms

The first step of the WFbC method involved clustering the DS waveforms into two groups via GMM. The GMM input was composed of the DS waveform amplitudes within a time window from −15 to 15 ms relative to the peak. Since the LFPs were sampled at 1 kHz, each DS waveform was represented by 31 samples (also referred to as features). This specific time window for feature selection yielded the highest mean accuracy among different window sizes for mouse A (see Results). For each channel, the mean DS classification accuracy was obtained across 20 GMM runs with different seeds for the initialization parameters.

The use of GMM was justified since most amplitude distributions of DS types demonstrated Gaussianity at each millisecond, as verified by two-sided Kolmogorov-Smirnov test. Of note, the distributions around 4 ms from the peak showed slight skewness, with the majority not fitting Gaussians (see Fig S2).

In cases where the LFP contains numerous artifacts, some may pass the DS detection process and be bundled together during GMM clustering. To address this issue, if a particular cluster comprised less than 5% of the total detected DSs, it was considered a potential artifact group based on empirical observations. In such cases, a new round of GMM clustering was performed excluding this group. This iterative procedure continued until the GMM produced two groups whose DS percentages were between 5 and 95%.

The DS-type assignment was based on the difference between the mean waveforms following the peak. For this, we compared the sum of the mean waveforms from 10 to 50 ms after the peak. The DS2 group was identified as the cluster with the lowest sum since the DS2 mean waveform (derived from CSDbC) exhibited more pronounced negative deflections after the peak.

A last and optional step involved verifying whether the DS clusters were actually of the same type. This was achieved by analyzing the second derivatives of their mean waveforms, as described below.

### DS single type checking

To assess whether the two DS groups, classified by either of the classification methods, represented distinct populations, we calculated a dissimilarity index (DI) based on the second derivative of their mean waveforms. Specifically, the DI was defined as the mean absolute difference between their scaled second derivatives (lying between 0 and 1) within a 20-ms window centered at the DS peak. If DI was equal to or lower than the empirically chosen threshold of 0.06 (adjusted to 0.07 in very few specific cases), it indicated that the DSs belonged to the same type.

The type of the single DS cluster was then determined through a voting process based on the width metrics and their corresponding boundary values. Type 1 was voted on if the DS width exceeded 19 ms, if the start width limit occurred before −9.5 ms from the peak, or if the end width limit occurred after 9.5 ms. Otherwise, votes were for type 2. In this way, DSs were classified as the type that received at least two votes out of three.

### Classification analysis

To evaluate the WFbC performance, we measured the classification accuracy as the percentage of DSs classified identically by the WFbC and CSDbC methods. Furthermore, we calculated the precision and recall metrics for each DS type. Of note, the repeated application of WFbC to a same dataset provided identical classifications for different GMM seeds (not shown).

The precision metric represents the fraction of correctly classified DSs of a specific type among all DSs classified as that type. The recall metric, on the other hand, measures the fraction of correctly classified DSs of a specific type out of all DSs that should have been classified as that type.

The confusion matrices are presented in their normalized versions with the values on their main diagonals corresponding to the recalls for DS1 and DS2.

### Statistical analysis

The coefficient of variation (CV), which is the ratio of the standard deviation to the mean, was used to contrast the spread of the DS metrics (M1-4) across channels and animals.

We performed two-tailed statistical hypothesis tests to compare medians considering a significance level of 0.05. The Mann-Whitney *U* test compared: the recalls and precisions between DS1 and DS2; and the scaled peak amplitudes and widths of DSs between genotypes for each DS type and age group and between the WFbC validation and application datasets. The Wilcoxon signed-rank test compared the paired values of the DS width metrics between the classification methods. The Kruskal-Wallis *H* test compared the DS widths across the three age groups of the same genotype. T-tests and Cohen’s D size effect were used to compare the mean waveforms between the well- and misclassified DSs by WFbC across both types.

Pearson’s correlation coefficient (*r*) measured the linear correlation between the metrics of DSs classified by both methods (CSDbC vs. WFbC) as well as between DS2 proportion and WFbC accuracy. The Wald test with t-distribution verified if the slopes of the linear regressions of the DS metrics returned by the classification methods were equal to 1.

### Software and algorithms

The data analysis was performed using custom-written codes in Python 3.8.11 with the following scientific packages: EMD 0.5.5 [37], Matplotlib 3.4.3 [38], NumPy 1.21.2 [39], Seaborn 0.11.2 [40], SciPy 1.7.1 [41], and Scikit-learn 0.24.2 [42].

## Results

### DS detection with artifact mitigation

To ensure the availability of reliable DS waveforms for classification purposes, we developed an accurate method for detecting DSs that goes beyond simply identifying LFP peaks above a specific threshold.

While the linear probe is essential for capturing laminar profiles needed for a precise DS classification [15], the detection of DSs requires just one channel. The electrode located at the hilus or the closest position was selected since DSs are generally most prominent in this particular DG layer [15]. In the case of mouse A, in which the probe crossed the DG, we considered one of the two linear arrays (red column in Fig 1A) to draw the laminar profile. The ninth electrode yielded the highest number of DSs, suggesting that it is located in the hilar layer. Furthermore, the analysis of CSD profiles during the occurrence of DSs (shown later) helped to estimate the probe position, as illustrated in Fig 1A.

DSs, being large and brief field events, can often be mistaken for externally generated patterns, such as abrupt deflections that affect all recording sites, referred to as artifacts. To address this issue, the DS detection process developed here contains optional steps for artifact mitigation (Fig 1B). The first step concerns the subtraction of the LFP of the target channel by the signal of a reference channel to eliminate common peaks. Here we took the apical channel of the linear probes as the reference. The resulting signal is then wideband filtered (1-200 Hz) to detect DSs as peaks above a certain threshold, as exemplified in Fig 1C. To avoid overlap between waveforms, the highest peaks are chosen to assure a minimum spacing between DSs. For more details, see the Materials and methods section.

Since the filtering process deforms the original signal, a peak offset correction was implemented to accurately represent the DS waveforms. Thus, each DS peak is determined as the maximum value of the raw LFP within a 20-ms window centered on the peak detected in the filtered LFP (Fig 1D).

For mouse A, approximately half of the 8098 peaks detected in the filtered LFP of channel 9 aligned with the maximum values in the raw LFP, thus no offset was applied. For most of the remaining half, an offset of −1 ms was required (Fig 1E). This adjustment resulted in an average DS waveform with a more pronounced peak compared to the waveform without the correction (Fig 1F).

Finally, to exclude remaining artifacts, we identified and discarded outliers in the amplitude distribution of the detected peaks. Outliers were defined as peak amplitudes that fell outside Tukey’s fences. For instance, clear artifacts were identified as outliers in channel 12 of mouse A, as can be observed in Fig 1G-J. Negative peaks may arise as a result of the detection process occurring after filtering. I.e., the peak appears positive in the filtered signal, but negative in the raw signal due to a very slow wave. Although this procedure is optional as it may also ignore valid DSs, we applied it to all data to prioritize identifying genuine DSs over slightly higher mean peak amplitudes and DS rates.

### DS rate is correlated to the power of delta oscillations

Following the detection of DSs, we investigated the relationship between their occurrence and the brain state, which is known to correlate with the behavioral state, as assessed indirectly through the relative power of delta-frequency oscillations.

As outlined in the Materials and methods section, we performed empirical mode decomposition on the LFPs of channels with evident DSs and focused on the fifth intrinsic mode function (IMF-5). The power spectral density (PSD) of IMF-5 consistently exhibited a pattern that matched the delta frequency band across all relevant channels, as demonstrated in Fig 2A and Fig S3. Therefore, we took IMF-5 to represent the LFP delta component, as exemplified in Fig 2B.

DSs were mainly observed during periods of increased delta power, as depicted in Fig 2C. These periods were defined here as “strong delta” when the relative delta power exceeded 0.9 (Fig 2D-F). Conversely, when the relative delta power was equal to or less than 0.9 (referred to as “weak delta”), the median rate of DSs was nearly zero in all relevant channels, as shown in Fig 2G (of note, despite the great rate difference, DS amplitude did not differ between weak and strong delta periods). Hence, when comparing DS rates in subsequent analyses, we specifically refer to DSs detected during periods of strong delta. Note in Fig S3A that the normalized mean PSDs of delta waves from mice A and B1 exhibit broader distributions, extending to higher frequencies. This was related to prolonged periods of weak delta activity in the LFP recordings of these specific animals.

With the assurance that the detected DSs were legitimate, supported by previous findings on the predominant occurrence of DSs at behavioral states characterized by increased delta power [15,16], we proceeded to analyze the waveforms of the DS types classified according to their laminar profiles.

### DSs can be classified as types 1 and 2 based on their laminar profiles

DSs are categorized into types 1 and 2 according to their origin, whether they arise from excitatory discharges originating in the LEC or the MEC, respectively. To determine the DS type, we analyzed the CSD profiles generated during the DS peaks using the method referred to as CSDbC. This method served as the reference for evaluating the performance of the WFbC method developed in this study, which classifies DSs from their own waveforms.

Fig 3A provides examples of DS1 and DS2 captured during a 1-second window from the recording session of mouse A. Consistent with previous observations [15], DS1 propagates across a greater number of channels on the linear probe compared to DS2. This is further supported by the laminar profiles depicted in Fig 3B, which display the mean peak amplitudes for each DS type. Additionally, the concatenated CSDs presented in Fig 3C demonstrate regular patterns of current sinks and sources along the DS types and, therefore, a consistent classification. Based on these findings, we can infer that DS1 current sinks correspond to the outer molecular layers of the DG (targeted by LEC afferents), while DS2 current sinks align with the middle molecular layers (targeted by MEC afferents). By combining this information with the understanding that the largest DSs are located in the hilus, we were able to estimate the respective layers associated with each channel, as illustrated in Fig 3B.

**Fig 3.**
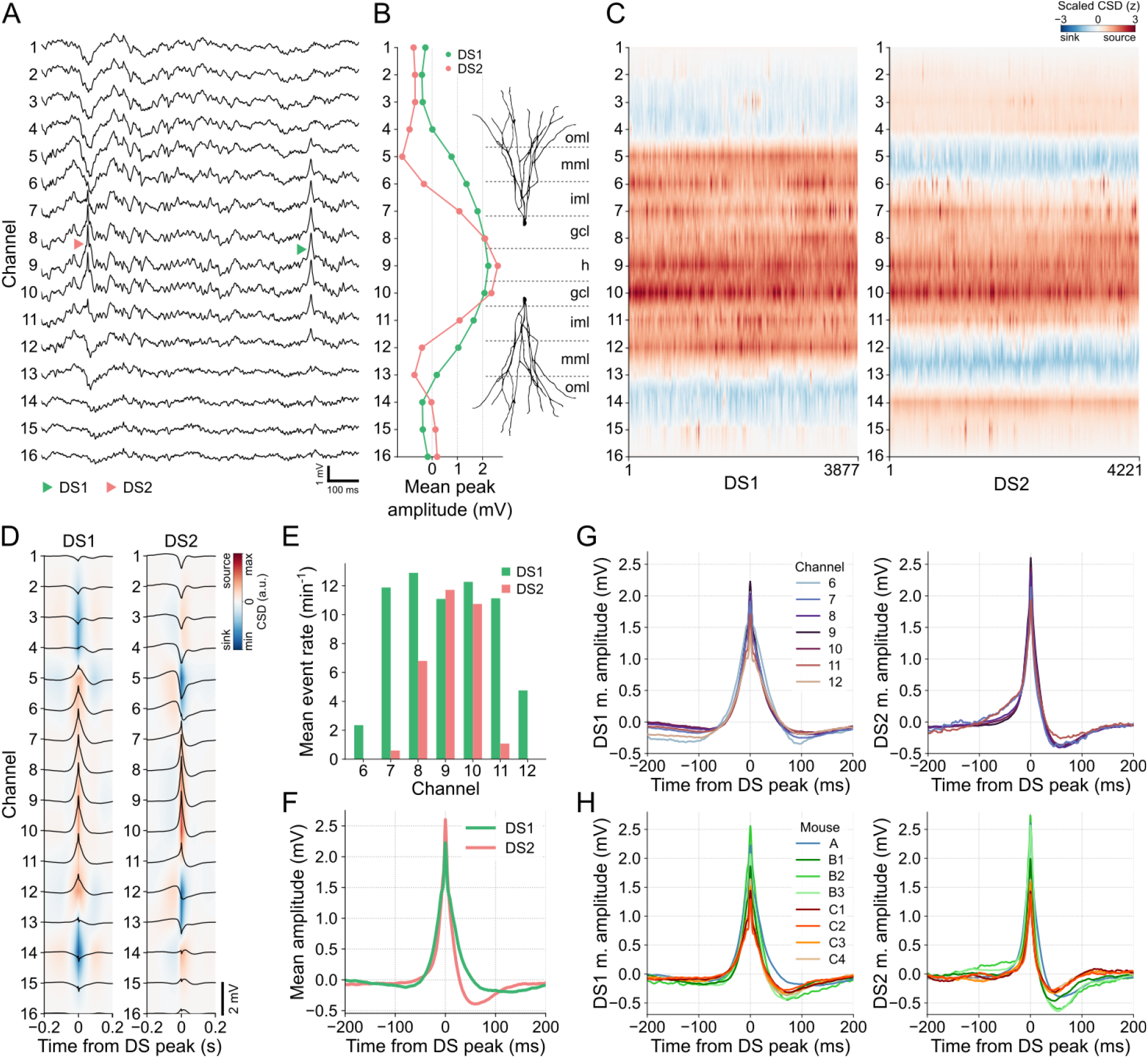
DSs can be classified as types 1 and 2 based on their laminar profiles. **(A)** Examples of each DS type in the raw LFP of mouse A. DS1 (green triangle) extends from channels 6 to 12, while DS2 (red triangle) is restricted to channels 7 to 11. **(B)** Mean peak amplitudes of each DS type across channels and estimated DG layers. **(C)** Concatenated CSDs, triggered at the DS peaks, for DS1 and DS2. **(D)** Mean CSDs of each DS type. Black traces are the mean waveforms in each channel. **(E)** Mean rate of DS types in each relevant channel. **(F)** Mean waveforms of each DS type in channel 9. **(G)** The left panel shows the DS1 mean waveforms in channels 6 to 12. The right panel shows the same, but for DS2 in channels 7 to 11. **(H)** The left panel shows the DS1 mean waveforms in the channel with more DSs detected in each mouse. The right panel shows the same, but for DS2. CSD: current source density; DS1: dentate spike type 1; dentate spike type 2; gcl: granule cell layer; h: hilus; iml: inner molecular layer; mml: middle molecular layer; oml: outer molecular layer.

In Fig 3D, we can observe the average CSDs generated by each DS type, as well as the mean waveforms captured within a 400-ms window centered at the peak for DSs detected in channel 9. These visualizations provide valuable information about the spatial distribution of DSs. Specifically, DS1 propagates between channels 6 and 12, while DS2 concentrates between channels 7 and 11. This spatial pattern is further supported by the mean event rates across channels, as depicted in Fig 3E, where DS2 did not appear during DS detection in the extreme channels 6 and 12. Consequently, channels 6-12 for DS1 and 7-11 for DS2 are the relevant channels for subsequent analysis. It is worth noting that a larger number of DS1 were detected on channels 8 and 10, which correspond to the putative granule cell layers in the supra- and infra-pyramidal blades. This is because the detection thresholds in these layers are lower than in the hilus at the same time that DS2 is attenuated faster than DS1 along their radial propagations.

The mean waveforms of DS types exhibit distinct characteristics, as illustrated in Fig 3F for DSs detected in channel 9 (hilus). The DS2 waveform was higher and sharper compared to DS1. Furthermore, DS2 shows a more pronounced negative post-peak potential and faster recovery to the baseline state. These characteristics were consistently observed across channels (Fig 3G) and mice (Fig 3H), suggesting that waveform features can be adopted for the classification of DSs using a single electrode.

When comparing DSs across animals, a challenge arises in determining the DG layer where the electrode detecting the DSs is positioned. As observed, both the mean amplitude and rate of DSs can vary depending on location. To address this issue before classifying DSs by their waveforms, we explored alternative metrics that exhibit greater stability and consistency across locations.

### Stability of DS metrics across layers and animals

In addition to the amplitude and rate metrics, we also explored the width as a feature for comparing DSs. Although the width is commonly measured as the DS duration at the base or at the half-peak [15,27,30], these approaches rely on the amplitude, which varies with the electrode position. To overcome this limitation, we propose an alternative width measure based on the DS waveform dynamics, since the LFP shape is much less distorted than attenuated along its propagation through the extracellular medium [43].

As explained in the Materials and methods section, the DS width proposed here corresponds to the distance between the second most prominent upward concavities around the peak of the mean waveform. This is a measure computed from the average DS waveform since individual DS waveforms can reflect the dynamics of other overlapping events.

The upward concavities of the mean DS waveform correspond to positive values in its second derivative, as indicated by the black dots in Fig 4A,B for the DSs detected in channel 9 of mouse A. The most pronounced concavities were found near the peak, being confined within a 10-ms window surrounding it. Additionally, we observed that the next prominent concavities fell within the time bands highlighted in yellow in Fig 4A,B. Thus, to measure the width of the waveform, we defined the start limit as the local maximum of the second derivative between −15 and −5 ms relative to the peak. Similarly, the end limit corresponded to the local maximum between 5 and 15 ms.

**Fig 4.**
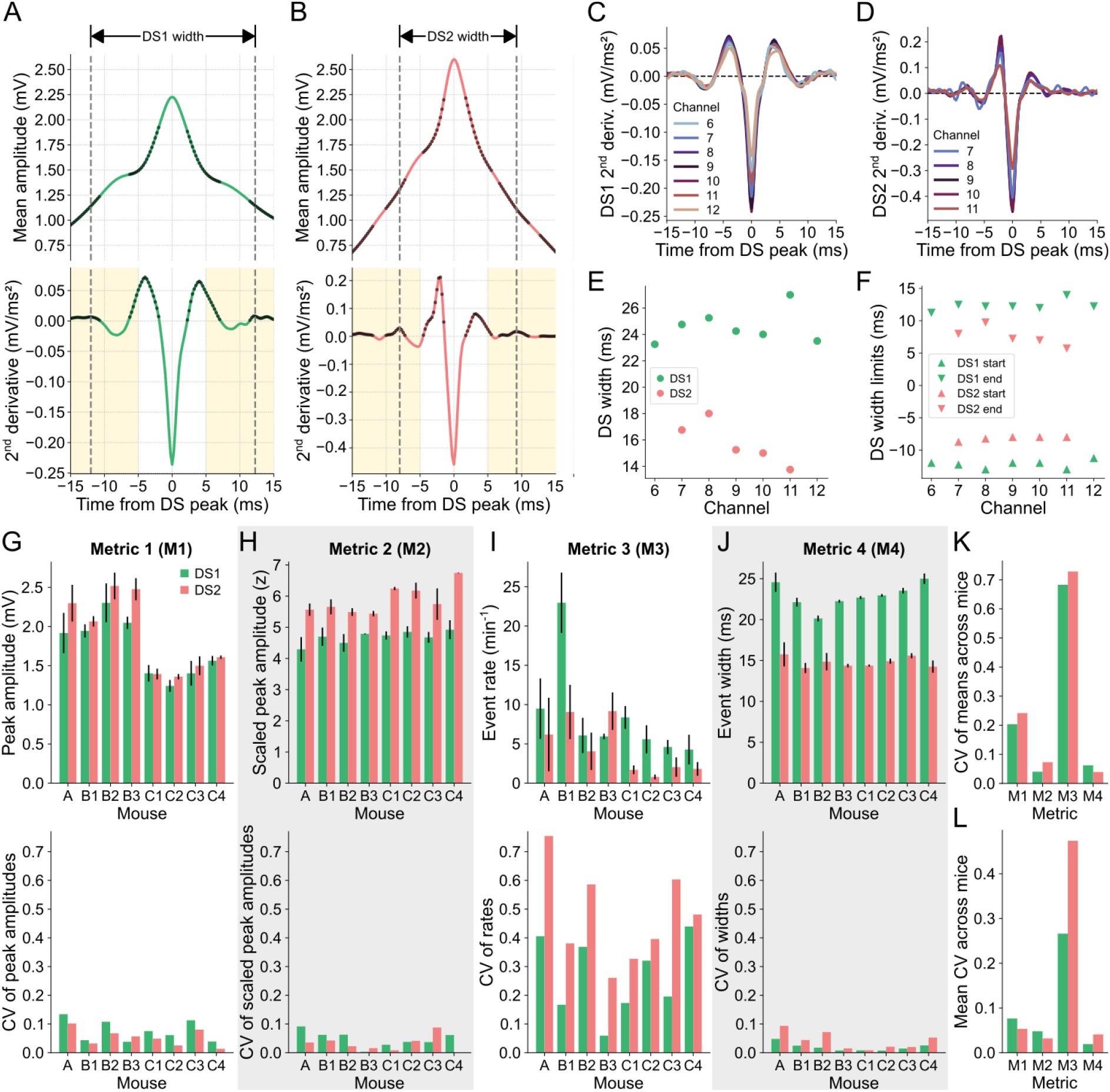
Stability of DS metrics across layers and animals. **(A)** DS1 mean waveform (top panel) in channel 9 of mouse A and its 2^nd^ derivative (bottom panel). Black dots mark the positive samples of the 2^nd^ derivative signal, which indicate the upward concavities of the waveform. Yellow shadows mark the range for the possible start (–15 to –5 ms) and end (5 to 15 ms) limits of the DS1 mean width. Dashed lines indicate the start and end limits of the DS1 mean width. **(B)** Same as A, but for DS2. **(C)** 2^nd^ derivatives of the DS1 mean waveforms in channels 6 to 12. **(D)** Same as C, but for DS2 in channels 7 to 11. **(E)** Mean widths of each DS type in each channel. **(F)** Mean width limits of each DS type in each channel. **(G)** The top panel shows the rates for each DS type (Metric 1 – M1) across channels for each mouse. Colored bars correspond to the means, and the error bars indicate the standard deviations. The bottom panel shows the coefficient of variation of the corresponding measurements in the panel above. **(H-J)** Same as G, but for the peak amplitudes (Metric 2 – M2) (H), the scaled peak amplitudes (Metric 3 – M3) (I), and the widths of each DS type (Metric 4 – M4) (J). Gray backgrounds indicate the metrics with the lower CVs. **(K)** Coefficients of variation of the means of each metric across mice. **(L)** Mean coefficients of variation of each metric across mice. CV: coefficient of variation; DS: dentate spike; DS1: dentate spike type 1; DS2: dentate spike type 2.

When detected on the other relevant channels, both DS types displayed regular profiles of the second derivatives of their mean waveforms, albeit with some noise observed in channels with fewer DSs, as shown in Fig 4C,D. The width values of both types of DSs detected in the relevant channels of mouse A are shown in Fig 4E. This animal exhibited the highest variability across channels, as depicted in Fig 4J. The DS width limits across channels, which are remarkably distinct between DS types, are shown in Fig 4F. In this way, the smaller width of DS2 is a reflection of the faster rise and fall of this type of DS, a conclusion also supported by the higher absolute value of the second derivative at the peak moment (Fig 4A,B). Of note, the mean waveform dynamics of each DS type were similar across mice (Fig S4).

To assess the variability of DS metrics across channels and mice, we employed the coefficient of variation (CV). The mean peak amplitudes and their standard deviations for both DS types across the channels of each mouse are presented in the top panel of Fig 4G, while the corresponding CVs are shown in the bottom panel. To enhance the stability of the amplitude metric, we z-scored the DS waveform amplitudes within a 400-ms window centered at the peak. Although the scaled peak amplitudes exhibited slightly smaller CVs on average across mice (Fig 4L), the CV values of the means across mice were considerably lower for both DS types (Fig 4K). The same analysis was applied to the event rate and width, as depicted in Fig 4I,J. Among all the DS metrics, the scaled peak amplitude and the width emerged as the most stable across channels and mice since they had the lowest CV of the means (Fig 4K) and lowest mean CV (Fig 4L) across mice. Of note, the width metric proposed by us also proved to be more stable than the width calculated at half height (Fig S5).

In possession of reliable and accurately classified DSs, along with suitable metrics for comparing them between experimental groups, we proceeded to the main goal of this study: the development of a classification method for DSs without relying on laminar profiles.

### DS classification based on waveforms

The WFbC method basically consists of two steps: DS clustering followed by the classification itself, i.e., the DS type assignment. Optionally, a final step can be applied to verify whether the groups represent distinct DS types. The macro flowchart of the method is depicted in Fig 5A.

**Fig 5.**
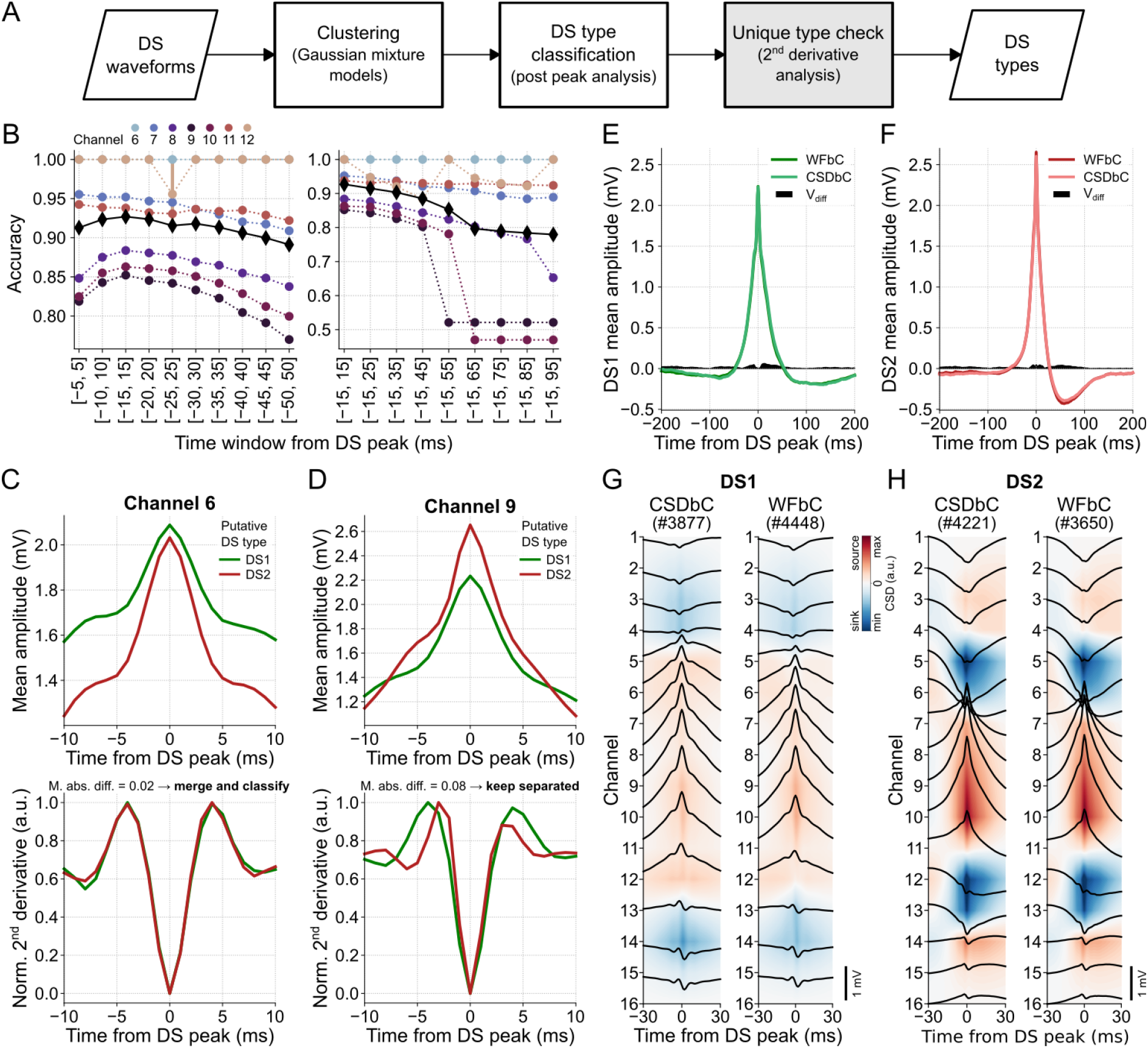
DS classification based on waveforms. **(A)** WFbC macro flowchart. After Gaussian mixture models (GMM) sort the DS waveforms into two clusters, they are classified as DS1 and DS2 by the post-peak analysis of the mean waveforms. Finally, the second derivative analysis checks if the two clusters represent a unique DS type, which will be classified according to its width. This step (in gray) can be bypassed. **(B)** Mean WFbC accuracies for different time windows and channels of mouse A across 20 different seeds of GMM initialization parameters. In the left panel, the time window increases by 5 ms before and after the DS peak. In the right panel, the time window increases by 10 ms after the DS peak. Circles and error bars indicate means and standard deviations for each channel. Black diamonds are the mean accuracies across channels. Overall, the [–15, 15] ms time window yields higher classification accuracy. **(C)** On channel 6, the 2^nd^ derivative analysis (bottom panel) shows that the two DS groups returned by previous steps of the method (top panel), in fact, correspond to a single group that will be classified as DS1 or DS2. **(D)** Same as C, but for channel 9. In this case, the 2^nd^ derivative analysis keeps the two groups separated. **(E)** Mean DS1 waveforms of channel 9 obtained by the CSDbC and WFbC methods. The absolute difference between them is shown by V_diff_. **(F)** Same as E, but for DS2. In E and F, V_diff_ < 0.07 mV. **(G)** Mean CSDs of DS1 classified by CSDbC (left panel) and WFbC (right panel) methods. **(H)** Same as G, but for DS2. CSD: current source density; CSDbC: CSD-based classification; DS: dentate spike; DS1: dentate spike type 1; DS2: dentate spike type 2; WFbC: waveform-based classification.

The clustering process used a GMM to divide the DS events into two distinct groups. Put simply, the GMM estimated the DS waveform distribution by assuming that they come from a combination of two different multidimensional Gaussians. Each Gaussian represented a cluster within the data, and the model determined the likelihood of each DS instance belonging to each cluster.

Each DS instance comprised the waveform amplitude values within a 30-ms window around the peak. This specific window size provided the highest mean classification accuracy across channels among several window sizes tested, as demonstrated in Fig 5B. First, we conducted an analysis of various time windows centered around the peak, ranging from [−5,5] to [−50,50] ms with a step size of 10 ms. We evaluated the classification performance for each window size and selected the optimal window size that yielded the highest accuracy. Subsequently, we investigated the influence of the post-peak period by increasing the window size by 10 ms after the peak. Interestingly, the optimal window size for classification covered the DS width values, being slightly larger than the widest observed width (27 ms). Furthermore, the different GMM seeds presented stable classification performances, except in the case of DSs detected on channel 12 through the [−25, 25]-ms window.

Using a window size of 30 ms and a sampling rate of 1 kHz, we obtained 31 amplitude samples per DS waveform. To streamline processing, we explored dimensionality reduction techniques, such as Principal Component Analysis, on the same set of features. However, these attempts yielded lower classification accuracies. Similar results were observed when standardizing the input data or using the values of the second derivative as DS features (not shown).

The final step, referred to as “unique type check”, verified the similarity between the normalized second derivatives of the mean waveforms of each DS group. If the mean absolute difference between them, within a 20-ms window centered at the peak, was equal to or lower than a predefined threshold (see Materials and methods), the DSs were considered to be of the same type. In such cases, a new classification based on the DS width was performed to assign the DS type to which the group belonged. On the other hand, if the result exceeded the threshold, the DSs remained classified into groups of different types. For instance, all DSs detected on channel 6 of mouse A were DS1 since DS2 did not propagate to the related DG layer. Despite initially being clustered into two groups by GMM, both putative DS types exhibited similar dynamics, as indicated by the second derivatives of their mean waveforms in Fig 5C. In contrast, DSs detected on channel 9, for example, belonged to different groups according to the dissimilarity between their mean waveform second derivatives, as shown in Fig 5D.

The classification of DSs detected in channel 9 achieved an accuracy of 0.85, resulting in mean waveforms of DS1 and DS2 that closely matched those obtained using CSDbC. The absolute amplitude differences (V_diff_) between the related samples of the mean waveforms classified by both methods were below 0.07 mV, as depicted in Fig 5E,F. A detailed comparison across the relevant channels of each mouse is shown in Fig S6-13. In general, the mean waveforms obtained through WFbC closely resembled those obtained using CSDbC. The differences between the mean waveforms were more pronounced in DS2 detected in channels located farther from the hilus. This was primarily due to the lower number of DS2 detections in those channels, despite the high classification accuracy. Moreover, the mean CSDs of each DS type were very similar between the classification methods, as shown in Fig 5G,H and Fig S14,15.

Most DSs misclassified by WFbC showed a high class probability (> 0.9) in the CSDbC GMM (Fig S16A), indicating that their CSD profiles did not differ from the well-classified DSs. This is reflected in Fig S16B,C, which shows similar CSD profiles between the well- and misclassified DSs of each type. On the other hand, by analyzing the mean waveforms (Fig S16D-F), we noticed a greater similarity between the waveforms of the well-classified DS1 and those of the misclassified DS2, with their end width limits coinciding. This implies that the average post-peak dynamics of a small subset of DS2 is similar to that of DS1. Interestingly, the misclassified DS2 were generated by weaker EC input, as shown in Figure S12C. Future research should delve into the relationship between EC discharge features and the DS dynamics.

### High performance of WFbC across animals

The accuracies achieved on channels with the highest number of DSs in each mouse ranged from 0.82 to 0.91. The normalized confusion matrices are shown in Figure 6A, where the main diagonal elements represent the percentages of correctly classified DSs for each type, i.e., the recall values. Notably, DS1 exhibited higher recalls than DS2 in all cases.

**Fig 6.**
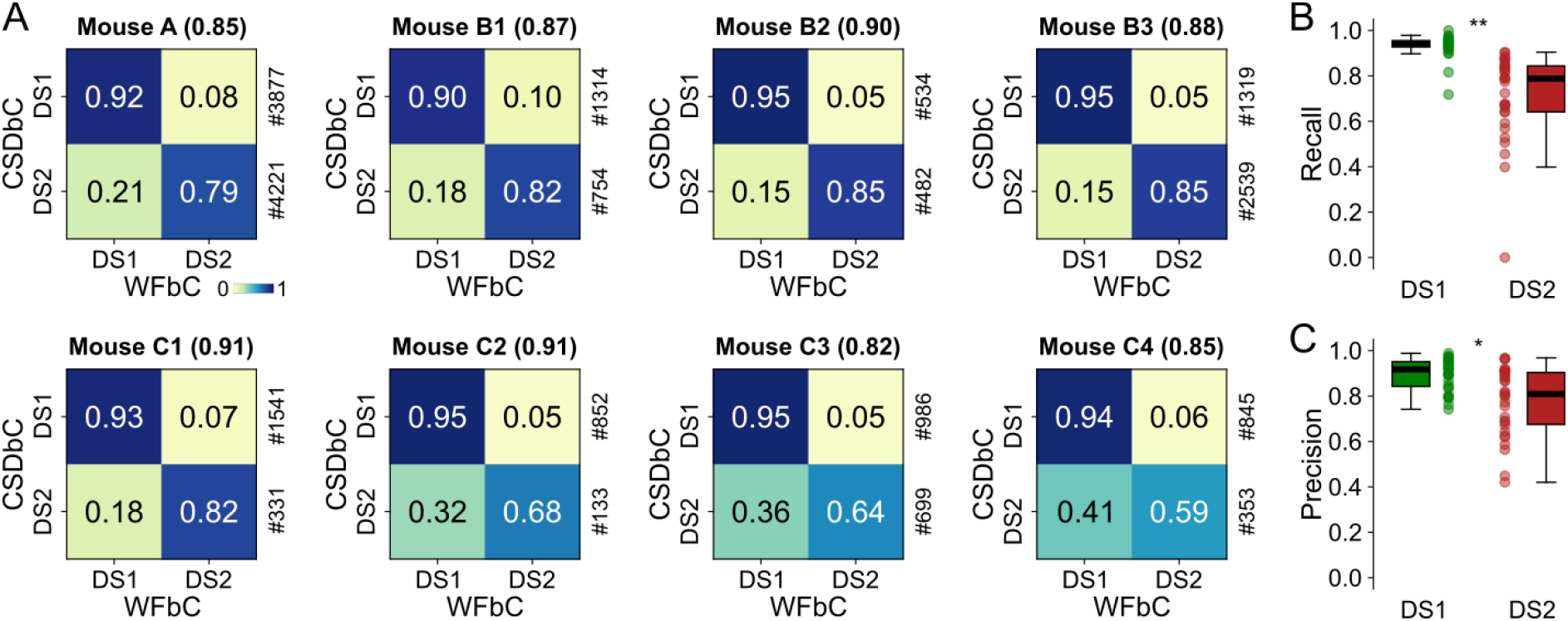
High performance of WFbC across animals. **(A)** Normalized confusion matrices for the channel with more DSs on each mouse. The classification accuracy is in parentheses, and the amount of each DS type returned by CSDbC is on the right. Note that the recall metric (main diagonal) is larger for DS1. **(B)** Boxplots show different recall distributions for DS1 and DS2 in all relevant channels of all mice. **(C)** Same as B, but for precision metric. n = 31 channels for each group in (B,C). CSD: current source density; CSDbC: CSD-based classification; DS1: dentate spike type 1; DS2: dentate spike type 2; WFbC: waveform-based classification. *p < 10^−3^, **p < 10^−9^ (Mann-Whitney).

When considering all relevant channels of all mice, the recall and precision distributions for DS1 were significantly higher compared to DS2 (Figure 6B,C), indicating that WFbC classifies DS2 more erroneously than DS1. On the other hand, the classification of DS1 is more reliable, especially when they constitute the majority. Thus, we further investigated the impact of DS-type proportions (percentages) on WFbC performance.

### WFbC performance depends on the balance between DS1 and DS2

WFbC tends to classify more DS2 as DS1 than the opposite, resulting in a lower DS2 proportion compared to CSDbC in most cases. Fig 7A shows a strong correlation between the DS2 proportions returned by both methods for the relevant channels of mouse A. Nevertheless, the slope of the linear regression significantly deviates from 1 (reference dashed line), as indicated by the *p*-value. Note that the points corresponding to channels 6 and 12 are overlapping on the zero value, anchoring the regression line. This implies that the difference in the number of DSs classified as type 2 between the methods becomes more pronounced as the DS2 proportion increases. Fig 7B shows similar results from the group data analysis, which considered the relevant electrodes from all animals.

**Fig 7.**
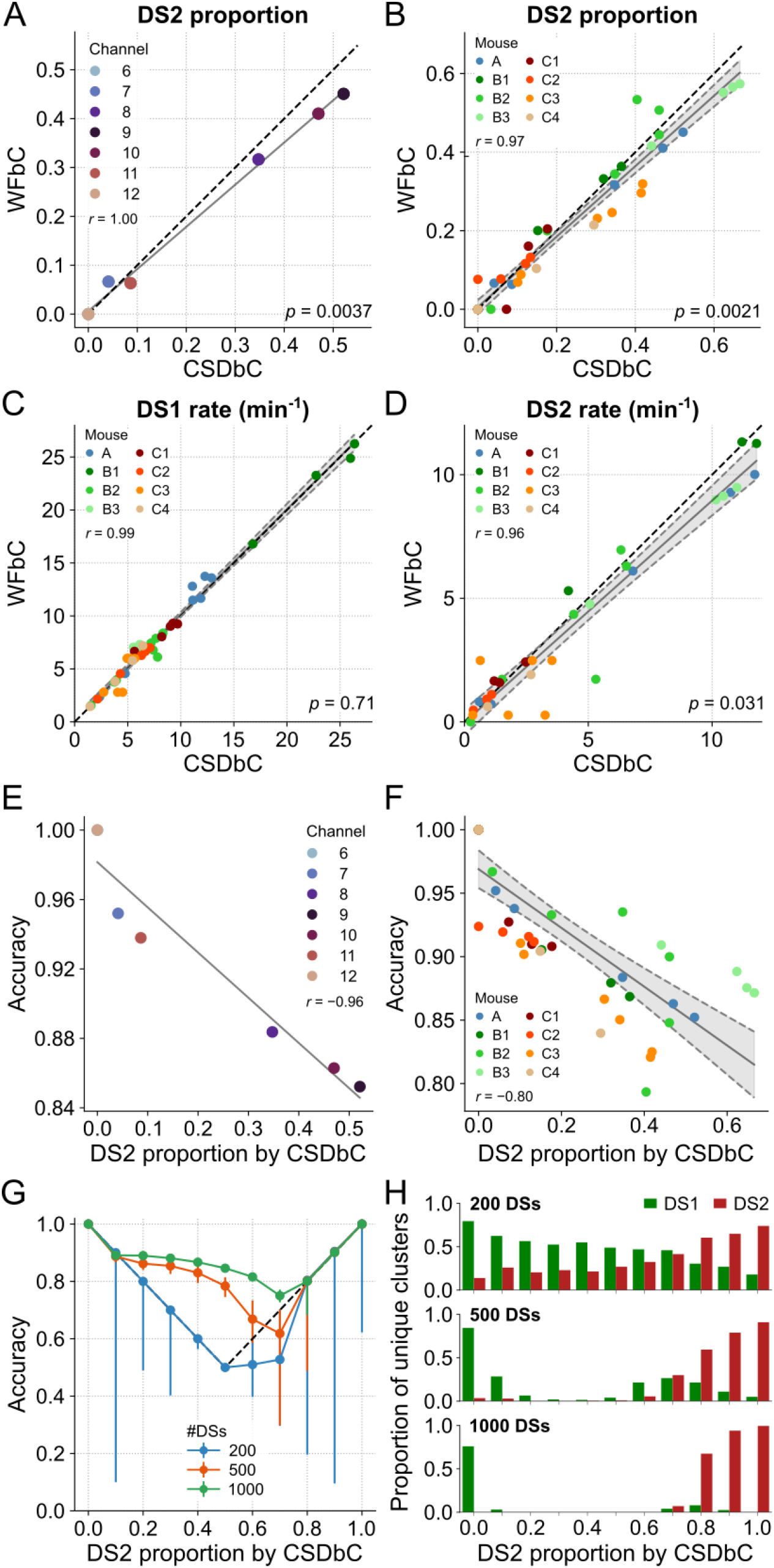
WFbC performance depends on the balance between DS1 and DS2. **(A)** DS2 proportions in each channel of mouse A returned by both classification methods. Despite the high correlation, the proportion of DS2 classified by WFbC is lower. The dashed black line denotes the y = x diagonal. Channels 6 and 12 have no DS2. **(B)** Same as A, but for all relevant channels of the 8 mice. Gray shadow represents the 95% confidence interval of the linear regression. **(C)** Same as B, but for DS1 rates returned by both classification methods. **(D)** Same as C, but for DS2. For A-D, the *p*-values refer to the significance of the slope against 1 (Wald). **(E)** WFbC performance across channels of mouse A is negatively correlated to DS2 proportion. **(F)** Same as E, but for all relevant channels of the 8 mice. **(G)** WFbC performance against DS2 proportion. Each dot is the median of 200 rounds with random and distinct DS choices from channel 9 of mouse A with different DS2 proportions. Error bars are the respective interquartile ranges. Black dashed lines indicate the classification chance, i.e. when WFbC returns a unique cluster: only DS1 when the DS2 proportion is less than 0.5, and only DS2 when the DS2 proportion is greater than 0.5. **(H)** The proportions of unique clusters returned by WFbC in G. The smaller the total number of DSs, the more unique clusters are falsely classified. CSD: current source density; CSDbC: CSD-based classification; DS: dentate spike; DS1: dentate spike type 1; DS2: dentate spike type 2; WFbC: waveform-based classification.

Since the DS-type balance relates to the rate, we next investigated the relationship between the rates of each type of DS classified by both methods. Fig 7C shows that the DS1 rates obtained by both methods for the relevant channels of all mice are strongly correlated. Importantly, the slope of the regression line did not significantly deviate from 1. However, this was not the case for DS2 rates, as shown in Fig 7D. On the other hand, the median absolute difference in DS2 rate between the methods was only 0.41 events/min, which is ∼14% of the DS2 median rate of 3 events/min returned by CSDbC. Along with the fact that the *p*-value was close to the significance level, we can conclude that the methods are consistent in determining both DS1 and DS2 rates.

Next, we hypothesized that the performance of WFbC would be inversely related to the proportion of DS2 due to its lower recall compared to DS1. Our suspicion was confirmed as the overall classification accuracy was higher for channels located farther from the hilus, where the DS2 proportion is lower. This relationship is clearly demonstrated in Fig 7E, which shows a strong correlation between the accuracy and DS2 proportion for the relevant channels of mouse A. The group analysis produced a similar result (Fig 7F), with accuracies consistently exceeding 0.8 and, thus, indicating an excellent performance of the WFbC method.

It is worth noting that channels on a probe that had smaller proportions of DS2 also had a lower total number of DSs detected. To further explore the relationship between WFbC accuracy, DS2 proportion, and the total number of DSs, we conducted a detailed investigation depicted in Fig 7G. Specifically, we examined the median accuracy over 200 classification rounds, randomly selecting DSs from channel 9 of mouse A for several DS2 proportions across three different total DS amounts: 200, 500, and 1000. Error bars indicate the interquartile ranges and the reference dashed lines denote the accuracy achieved if all DSs were classified as a single type: DS1 below 0.5; DS2 above 0.5; and DS1 or DS2 at 0.5.

As observed in the graph, the total number of DSs is critical for the effective performance of the WFbC method. The accuracy improves as the number of DSs increases. However, as mentioned earlier, the classification accuracy declines as the DS2 proportion rises. The magnitude of this decline is determined by the total number of DSs. With a high number of DSs, such as 1000, the accuracies remain above the reference values up to a DS2 proportion of 0.7. Above it, the median accuracy corresponds to the reference value. With a smaller number of instances, like 200 DSs, the WFbC tends to merge DSs into two pre-clustered groups and mistakenly categorize them as a single type, as shown in Fig 7H. Based on this analysis, it is recommended to have more than 500 DSs as input data for a reliable application of the WFbC method.

### Similar widths for DSs classified by either WFbC or CSDbC

Lastly, we examined the widths of each DS type across the classification methods. Fig 8A shows that the DS1 widths from the relevant channels of all mice exhibited a strong correlation, with no difference between the slopes of the linear regression and the reference line. On the other hand, the correlation for DS2 widths was lower, and there was a significant difference between the slopes. However, the width distributions were not statistically different, as indicated in Fig 8B, and the mean absolute difference between the methods was merely 0.29 ms for DS1 and 0.57 ms for DS2.

**Fig 8.**
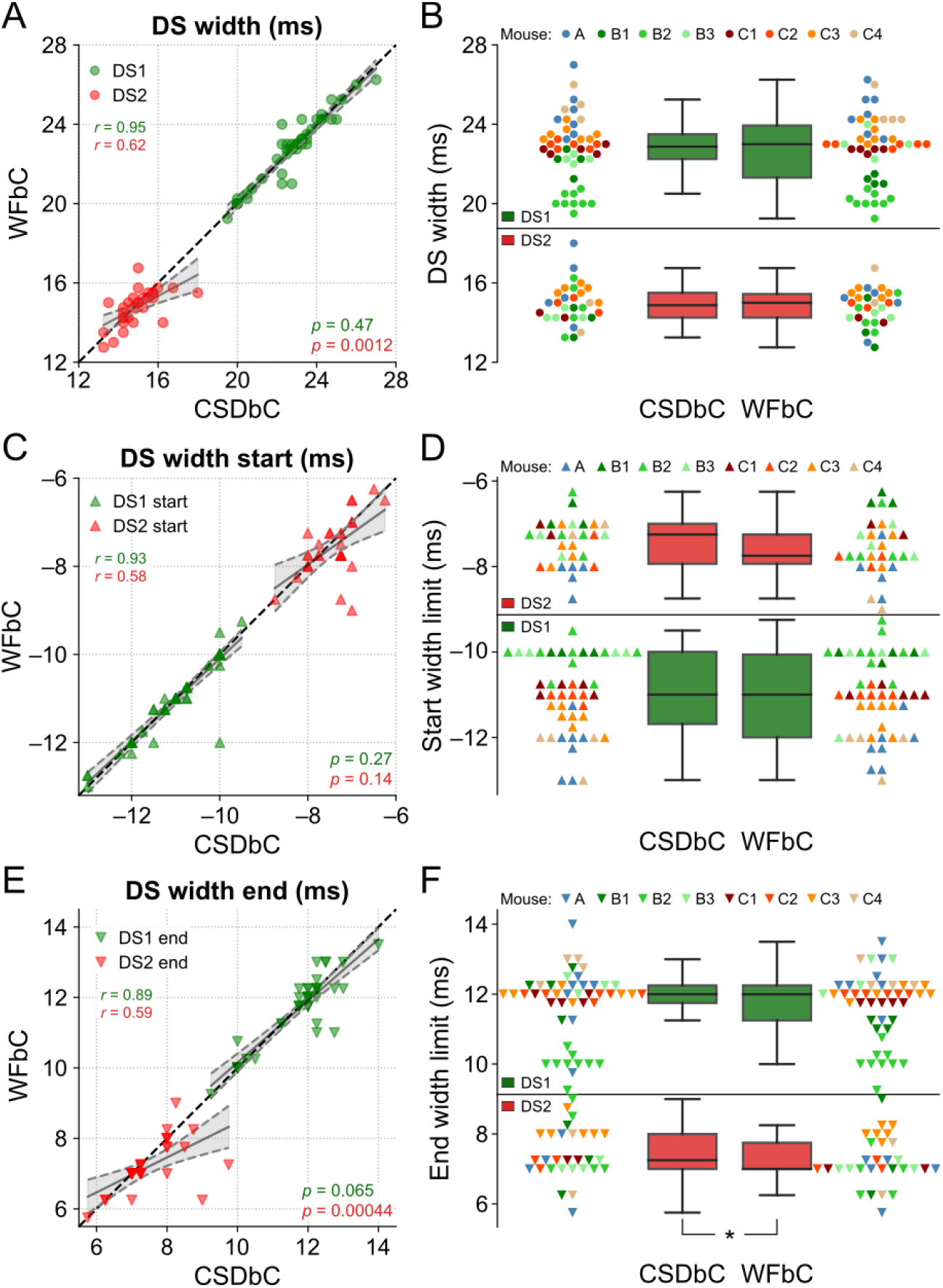
Similar widths for DSs classified by either WFbC or CSDbC. **(A)** Widths of each DS type for both methods. The dots correspond to the channels of the 8 mice. The dashed black line denotes the reference y = x diagonal. **(B)** Comparison of the width distributions of each DS type between methods. The dots of the same color correspond to the channels of the same mouse. There is no statistically significant difference in the comparisons of both DS types through the Wilcoxon signed-rank test. **(C,D)** Same as A and B, but for the start limits of the widths of each DS type. **(E,F)** Same as A and B, but for the end limits of the widths of each DS type. In A, C and E, the *p*-values refer to the significance of the slope against 1 (Wald). CSD: current source density; CSDbC: CSD-based classification; DS: dentate spike; DS1: dentate spike type 1; DS2: dentate spike type 2; WFbC: waveform-based classification. \**p* = 0.014 (Wilcoxon).

In Fig 8C-F, it is noticeable that the end width limits were responsible for the more pronounced difference between the DS2 widths. While there was no significant difference between the slopes of the linear regression and the reference line between the start width limits for both DS types, there were significant differences for the slopes referred to the end width limits of DS2 across methods and a slightly significant difference between their distributions, with *p*-value close to the significance level.

Taken together, these results provide evidence that the widths of the DS types obtained through WFbC are consistent with those classified by CSDbC, with a small bias associated with DS2.

### ApoE4-KI mice exhibit wider DSs compared to age-matched ApoE3-KI mice

As an application of the WFbC method, we investigated potential differences in the waveforms of both DS types between ApoE3-KI and ApoE4-KI mice. The latter, serving as an AD model, is known to experience early-onset impairment of hilar GABAergic interneurons [44–46].

The electrophysiological recordings in the dorsal hippocampus of each mouse were conducted through a 32-channel silicon probe with electrodes distributed across 4 shanks. The vertical distance between electrodes was 200 µm, thus not permitting the DS classification via CSD analysis. For each animal, we selected the DG channel that yielded more DSs with a minimum of 500 and a mean peak amplitude above 1 mV.

Following the DS classification via WFbC, we evaluated the scaled peak amplitudes and the widths of each DS type between genotypes within age-matched groups, and across ages within the same genotype. We chose these metrics since they demonstrated greater stability across DG layers, ensuring reliable comparisons among DSs detected on distinctly located electrodes.

At first glance, the mean waveforms of each DS type appear consistent across genotypes within the age-matched groups, as depicted in Fig S17,18. Curiously, a few adult and old mice had no DSs classified as type 1, indicating either the absence or a low occurrence of DS1 in the analyzed sessions. Thus, the DS2 comparisons include more animals.

In terms of the scaled peak amplitudes, we found no significant differences between the two genotypes within the age-matched groups for each DS type (Fig S19A-C). Conversely, the DS widths were significantly different in most of the comparisons, with the ApoE4-KI groups exhibiting wider waveforms (Fig 9A-D,G-J). The widths of DS1 in adult mice were the only case without a significant difference between genotypes. Overall, the width differences were more pronounced for DS2 than DS1, resulting in a longer recovery time to baseline activity in ApoE4-KI mice (Fig S18). This observation suggests potentially different levels of disease-related impairment in the LEC- and MEC-DG circuits. Importantly, the DS widths remained statistically similar across the three age groups within each genotype (Fig S19D,E), suggesting no significant effect attributed to the aging process.

**Fig 9.**
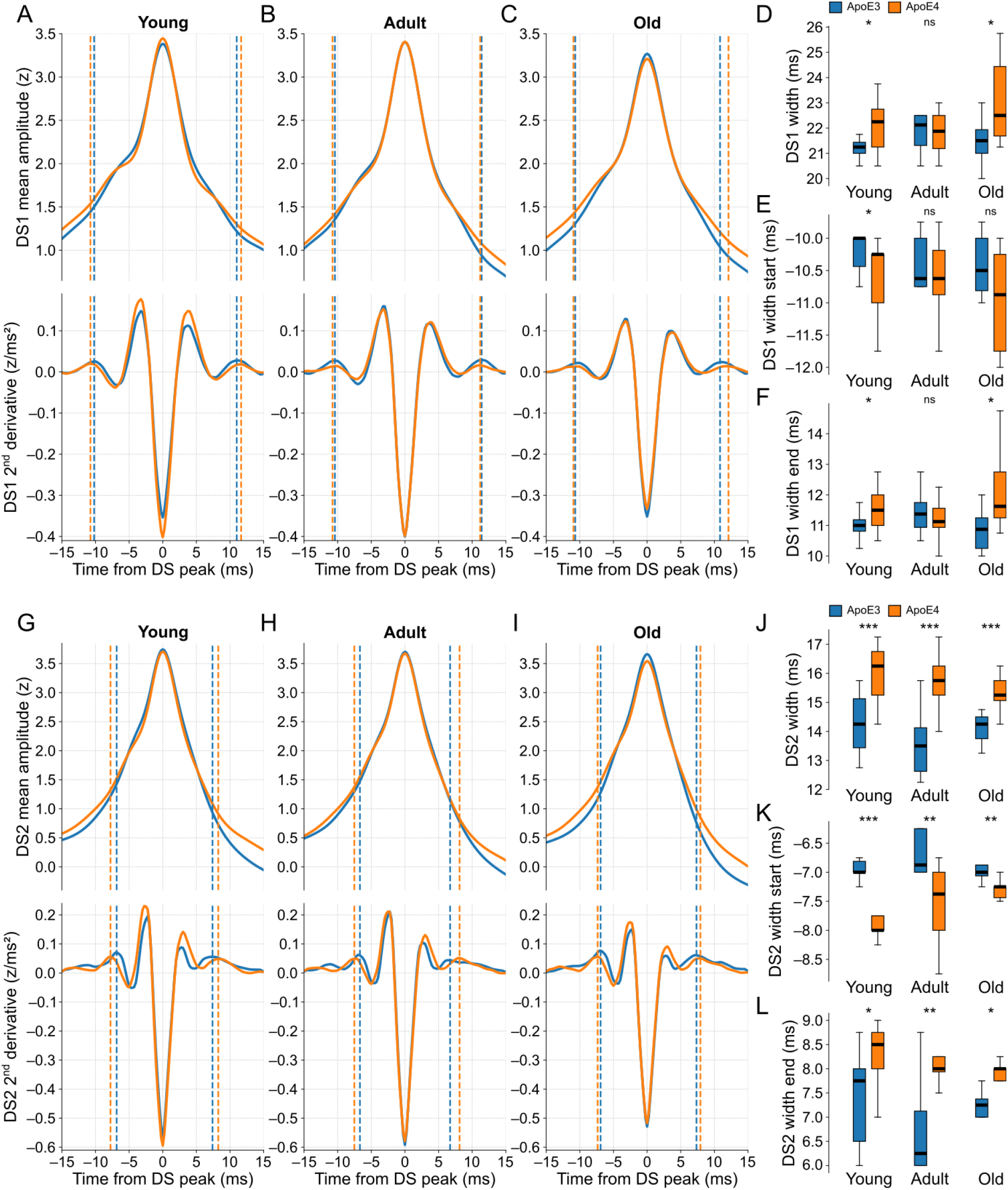
ApoE4-KI mice exhibit wider DSs compared to age-matched ApoE3-KI mice. **(A)** The top panel shows the scaled DS1 mean waveforms of ApoE3-KI (blue trace) and ApoE4-KI (orange trace) mice when young (n = 10 and 13). The bottom panel shows the respective second derivatives of the top signals. The dashed lines are the width limits, whose start and end limits are before and after the DS peak respectively. **(B)** Same as A, but for adult mice (n = 8 and 12). **(C)** Same as A, but for old mice (n = 8 and 10). **(D)** Boxplots comparing the DS1 width distributions between genotypes in each age group. **(E)** Same as D, but for the start limits of the DS1 widths. **(F)** Same as D, but for the end limits of the DS1 widths. **(G-L)** Same as A-F, respectively, but for DS2. DS types were classified by WFbC. ApoE3-KI: Apolipoprotein E3 knock-in mice; ApoE4-KI: Apolipoprotein E4 knock-in mice; DS1: dentate spike type 1; DS2: dentate spike type 2; ns: not significant. *p < 0.05, **p < 0.01, ***p < 0.001 (Mann-Whitney).

The observed differences in DS widths between genotypes were indicative of alterations in both pre- and post-peak dynamics, as evidenced by the variations in the distributions of the corresponding width limits. The modest differences in DS1 widths were reflected by similarly mild differences in the corresponding width limits (Fig 9D-F). In the case of DS2 widths (Fig 9J-L), particularly in the young age groups, the disparities between genotypes were more pronounced in the start limits. This result suggests that the neuronal mechanisms underlying the rise of DS2 may be affected before those responsible for its falling dynamics.

Although the scaled peak amplitudes between the ApoE mice (classified via WFbC) and those from datasets A-C (classified via CSDbC) differed significantly, their DS widths did not (Fig S19F,G). Beyond validating the WFbC method, this result reinforces the preservation of the dynamics of each DS type, as represented by the width measurements, across DG layers and mice cohorts. Therefore, this finding emphasizes the robustness and reliability of the width as a measure of the DS dynamics.

The broader DS waveforms observed in ApoE4-KI mice may indicate increased neuronal activity in the EC-DG circuit, consistent with previous findings in the hippocampal formation of this AD model [47]. Furthermore, these results demonstrate the effectiveness of WFbC in classifying DSs, allowing its use for expanding the research on their roles in memory processing.

## Discussion

DS types derive from excitatory discharges from different regions of the EC [15], each associated with specific content for the formation of episodic memories [48]. While LEC processes contextual information and selectively projects to the DG OML via the lateral perforant path (LPP) [49–52] – generating DS1 – inputs from MEC via the medial perforant path (MPP) convey spatial cues and specifically target the DG MML [49,53–55] – producing DS2. The distinguished involvement of MPP and LPP in memory processing is supported by their distinct ability to induce long-term potentiation [56], beyond the LPP preference for innervating newly integrated granule cells [57], which modulate entorhinal inputs into the DG [58]. Thus, we can gain insights into the specific contributions of DS types to cognitive processes by studying their properties and interactions within the hippocampal circuitry. However, since the first description of DS types three decades ago [15], little has been explored about their involvement in hippocampal-dependent cognitive functions [59,60]. A constraining factor for DS classification is the requirement of a laminar profile that discerns current sink positions within the DG molecular layer, which is only achieved through minimally spaced electrodes [15]. To overcome this limitation, we developed a waveform-based classification method that requires a single recording site. This novel approach extends the investigation of DSs by allowing, for instance, their classification in recording setups that capture a larger number of cells, like tetrodes [28], which provide additional features of the same spike for more accurate sorting [29].

The robust performance of WFbC indicates that the waveforms of each DS type carry information about their respective origins. This is consistent with electrophysiological evidence demonstrating distinct characteristics of excitatory postsynaptic potentials (EPSPs) elicited by the stimulation of LPP and MPP. Specifically, MPP stimulation induces EPSPs that are higher in amplitude and faster in onset compared to LPP stimulation [61], mirroring the differences observed between DS2 and DS1 waveforms. MPP stimulation also leads to greater amplitudes of population spikes, revealing a more effective activation of DG granule cells [61], consistent with more protruding current sinks and sources produced by DS2. Although there is a positive correlation between the rise time of EPSPs and the synaptic distance from the soma [62,63], these activation features have been attributed to active properties of synaptic transmission rather than differential passive decay of dendritic depolarization [64,65]. Another fact that seems to associate the divergent interactions between MEC and LEC inputs to differences in receptor make-up of granule cells is that MEC-DG synapses are predisposed to undergo potentiation, while LEC-DG synapses tend to express synaptic depression [66]. These observations may give WFbC an advantage in classifying DSs in species where the radial distribution of LPP and MPP is not well defined. Unlike rodents, the EC-DG projections of primates and bats are mixed in the MML and OML [67,68], theoretically making the classification of DSs unfeasible via CSD analysis.

One of the main challenges in investigating DSs is ensuring that legitimate events have been detected. Unlike oscillatory patterns, which are identified from spectral analysis, DSs are typically detected as peaks in the LFP greater than a certain threshold [16,18,19,24–27,69,70]. This makes DSs easily mistaken for artifacts or even prominent peaks of gamma waves, as we believe to have happened in a previous study [71]. Regarding the latter, the authors found high DS rates even during mouse mobility [71], contradicting the well-established DS predominance during quiet behavioral states and slow-wave sleep [15,16], when memory consolidation is thought to occur [72]. The events detected are further questioned due to the low peak amplitudes (∼0.6 mV) and the oscillatory patterns (∼45 Hz) in their mean waveforms [71]. Moreover, it is known that DSs and DG gamma peaks evoke similar CSD dipoles, with the power of the latter increasing during mobility and dramatically decreasing following perforant path ablation [73]. Taken together, these observations raise concerns about the reliability and validity of the detected events as true DSs in some previous studies, leading us to believe that they actually corresponded to prominent peaks of gamma waves.

Studies conducted in urethane-anesthetized rats have also shown DSs with low peak amplitudes and oscillatory waveform profiles [18,74], exhibiting inconsistent results with pre-existing literature. One contradiction concerns the synchronous occurrence of DSs between the two hemispheres. While Bragin and colleagues observed interhemispheric synchrony for most events [15], Lehtonen and colleagues recently described the opposite, advocating rodent hippocampus lateralization [74]. Other conflict concerns the mossy cell recruitment during DSs as described by Bragin and colleagues [15] and also by Senzai and Buzsáki [26]. This finding was contradicted by Sanchez-Aguilera and colleagues, who reported a DS silencing effect in this neuron type [18]. To resolve such discrepancies, future studies should consider the anesthetic state of the animals, along with the techniques used for DS detection and the network activity emerging from the different DS types.

Comparing the peak amplitude of DSs between different subjects is also tricky. Assuming that the same type of electrode is used to record DSs, variations in their mean amplitudes are primarily attributed to differences in electrode placement within the DG. As seen, DS peaks are typically higher between the granule cell layers and decrease towards the molecular layers. Even in the same layer, variations in the density of cell bodies may influence the DS magnitudes across animals [17]. Furthermore, the use of varied recording electrodes involves considering additional factors, as pointed out by extracellular recording models, which can be extrapolated to the analysis of the DS waveforms. These factors include the surface area of the recording site, the impedance of the electrode-tissue interface, and the presence of local electrical inhomogeneities such as encapsulation, edema, and coatings around the electrode shank [75–77].

To minimize the impact of confounding factors when comparing the peak amplitudes of DSs, we z-scored the mean waveform amplitudes within a 400-ms window around the peak. In fact, the scaled peak amplitude resulted in lower coefficients of variation across the DG layers and animals (and therefore across electrode layouts) when compared to the absolute peak amplitudes and event rates. The latter metric showed the most variation since the number of detected DSs depends on their height, which in turn varies along the DG lamina. Another metric that showed minimal variation was the DS width measure proposed in this study. Unlike conventional width measurements that rely on amplitude [15,27,30], our proposed metric takes into account the dynamics of the average waveform, specifically the temporal distance between the second upward concavities around the peak. Further investigations should explore the activation sequences among different cell types within the DG network to understand the dynamics revealed by the second derivative patterns of the mean waveforms, which are used to identify the concavities that indicate the DS width limits, and thus gain deeper insights into the DS underlying mechanisms in the hippocampal circuit.

By using the scaled peak amplitude and the width of the mean waveforms, we compared morphological features between the DS types classified by WFbC in the transgenic mice, which represented an AD model (ApoE4-KI) and the control group (ApoE3-KI) across different ages. Our results revealed no differences in the scaled peak amplitudes across genotypes or across the age groups of the same genotype. On the other hand, ApoE4-KI mice exhibited wider DS waveforms with a higher effect size observed in DS2 comparisons. Of note, the width values of the DS types of the ApoE3/4-KI mice were equivalent to those optimally classified by CSDbC (datasets A-C). These differences in DS widths between genotypes are consistent with the ApoE4-associated hyperactivity in the hippocampal formation mainly due to a major loss of GABAergic interneurons [45,78–80]. This is also consistent with hyperactivity found in MEC of these mice [47], which would affect DS2s specifically. Aligned with this finding, individuals with a genetic predisposition to AD exhibit early spatial disorientation symptoms that have been related to dysfunction in the firing of MEC grid cells [81].

The WFbC method has demonstrated high performance, yielding consistent results for the analyzed DS metrics. We observed that robust performance is achieved with a sufficient number of detected events (> 500) with a minimum average height (> 1 mV), which can be attained through ample recording time during resting states with a well-implanted electrode. Additionally, DS classification performed better for a higher proportion of DS1 as they were more accurately classified. Considering the balanced natural occurrence between DS types (to be investigated), this condition is likely to be satisfied, as DS2 propagation is more restricted along the DG radial axis. Thus, our method proves to be viable and is expected to facilitate the analysis of DSs in previously challenging recordings, expanding the investigation of DS-type functional roles.

## Conclusion

Our results provide compelling evidence for the robust performance of WFbC in classifying DSs. By utilizing our method, the exploration of DSs can be extended to a wider range of recording techniques, facilitating a more comprehensive understanding of their functional significance in mnemonic processes.

## Data availability statement

The original data used in this study, as well as the access links, are described in the Materials and methods section. The derived data identified as the waveforms of DSs detected in the channel located at the hilus of each animal are available at https://data.mrc.ox.ac.uk/waveform (DOI: <u>10.5287/ora-wrbzrbwpk</u>). The codes for the detection of DSs and their classification by the CSDbC and WFbC methods are available at https://github.com/tortlab/dentate-spikes (DOI: <u>10.5281/zenodo.10080866</u>).

## Author contributions

Author contributions based on the CRediT Taxonomy.

Conceptualization: RMMS and ABLT.

Data curation: RMMS, VLdS and EAAJ.

Formal analysis: RMMS.

Funding acquisition: ABLT.

Investigation: RMMS.

Methodology: RMMS.

Project administration: ABLT.

Resources: ABLT.

Software: RMMS.

Supervision: ABLT.

Validation: RMMS.

Visualization: RMMS.

Writing – original draft: RMMS.

Writing – review & editing: ABLT, VLdS, EAAJ, DD and YH.

## Acknowledgments

We thank César Rennó-Costa, Cleiton Aguiar, Diego Laplagne, Hindiael Belchior and Renan Moioli for engaging in critical discussions that enriched our study. We are also grateful to Daniela Moura for providing the dentate gyrus image used to illustrate the schematics in Fig 1A, and Yuta Senzai and György Buzsáki for making dataset A publicly available.

