## Supplementary Figures for "Waveform-based classification of dentate spikes"

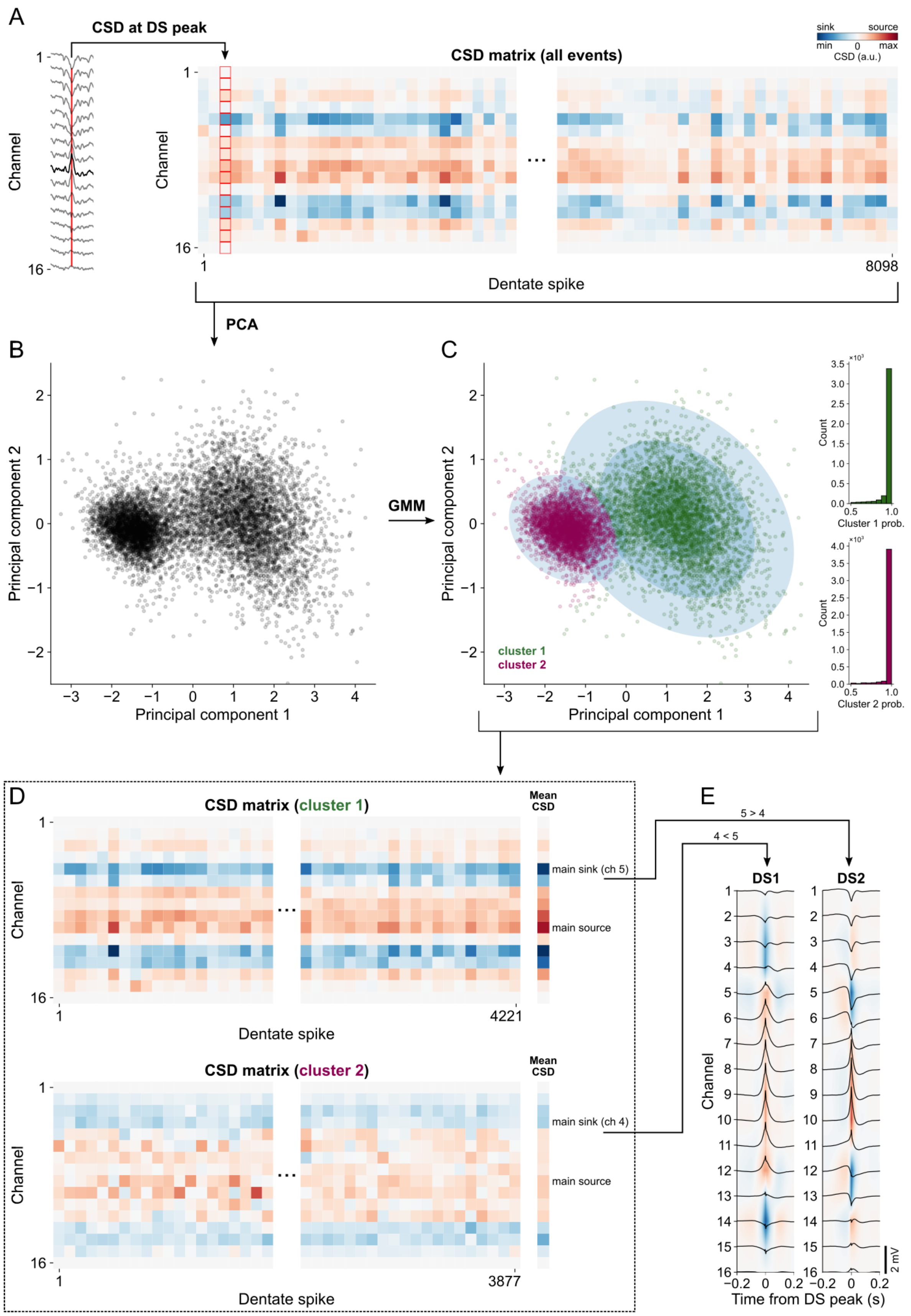


**Fig S1. Step by step of the CSDbC method using mouse A DSs.**

**(A)** After DS detection in the hilus channel (bold trace in the left panel), the CSD profiles are calculated at the peak of each event (red line), generating a CSD matrix of events by channels (right panel). **(B)** The CSD matrix is projected into the two dimensions of greatest covariance by PCA. **(C)** Events projected into the first principal component are clustered into two groups by GMM. The ellipses indicate the Gaussian probability distributions of each cluster in three levels. The right panels show high clustering probability for most events. **(D)** The CSD matrix of each cluster generates a mean CSD profile that indicates the position of the main (most prominent) current sink above the main current source. **(E)** DS type assignment for each cluster based on the positions of the main current sinks. Cluster 2 corresponds to DS1 since the main current sink is located in channel 4, which is above channel 5 (cluster 1 main sink). Panels show the mean CSD profiles and waveforms (black traces) for each DS type. ch: channel; CSD: current source density; DS: dentate spike; DS1: DS type 1; DS2: DS type 2; GMM: Gaussian Mixture Models; PCA: Principal Component Analysis.

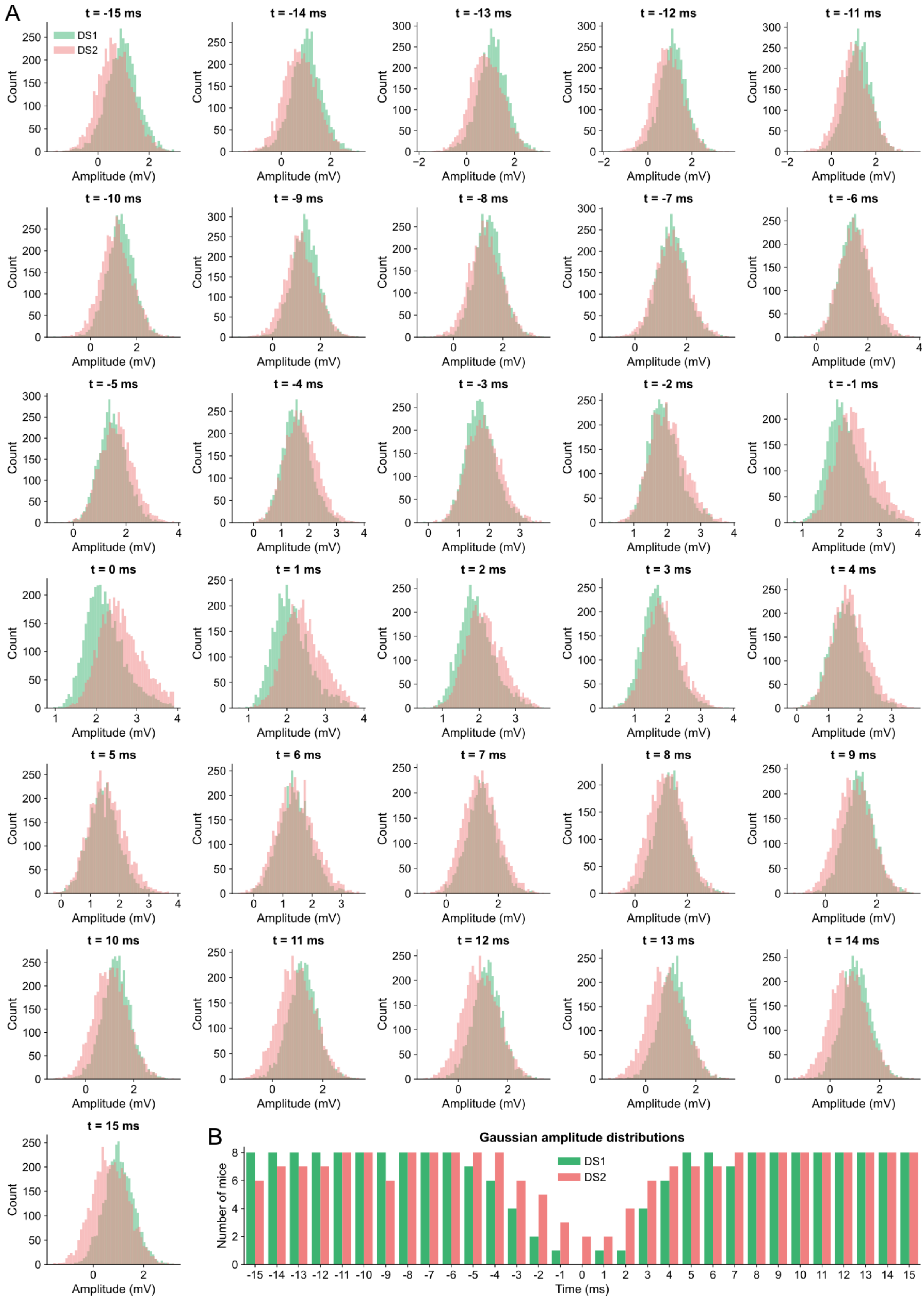


**Fig S2. Gaussianity of amplitude distributions of DS types.**

**(A)** Amplitude distributions of DS types in each millisecond within a 30-ms window around peak. **(B)** Number of mice whose amplitude distributions showed Gaussianity in each millisecond within a 30-ms window around peak.

DS1: dentate spike type 1; DS2: dentate spike type 2.

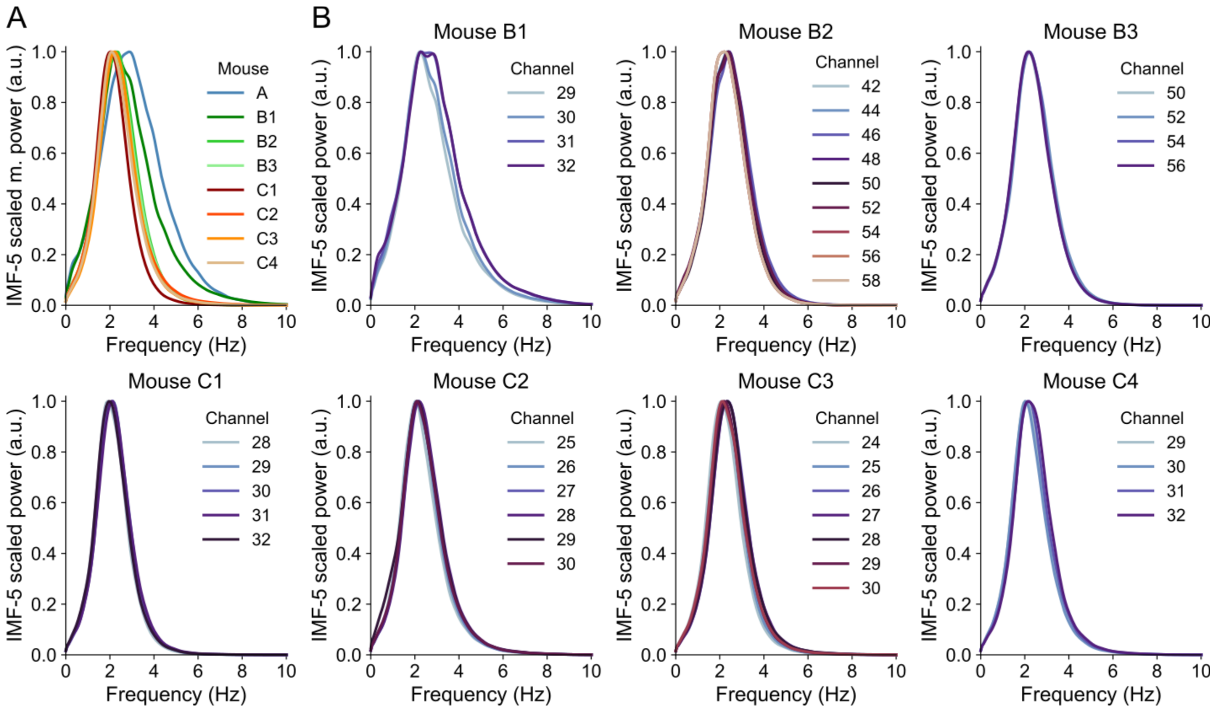


**Fig S3. Consistent PSDs of IMF-5 across mice and DG layers.**

**(A)** Normalized mean power of IMF-5 of all mice. Each trace is the mean PSD of one mouse across its relevant channels. **(B)** Each panel exhibits the normalized IMF-5 PSDs of all relevant channels of each mouse. The panel corresponding to mouse A is shown in the inset of Fig 2A. IMF: intrinsic mode function.

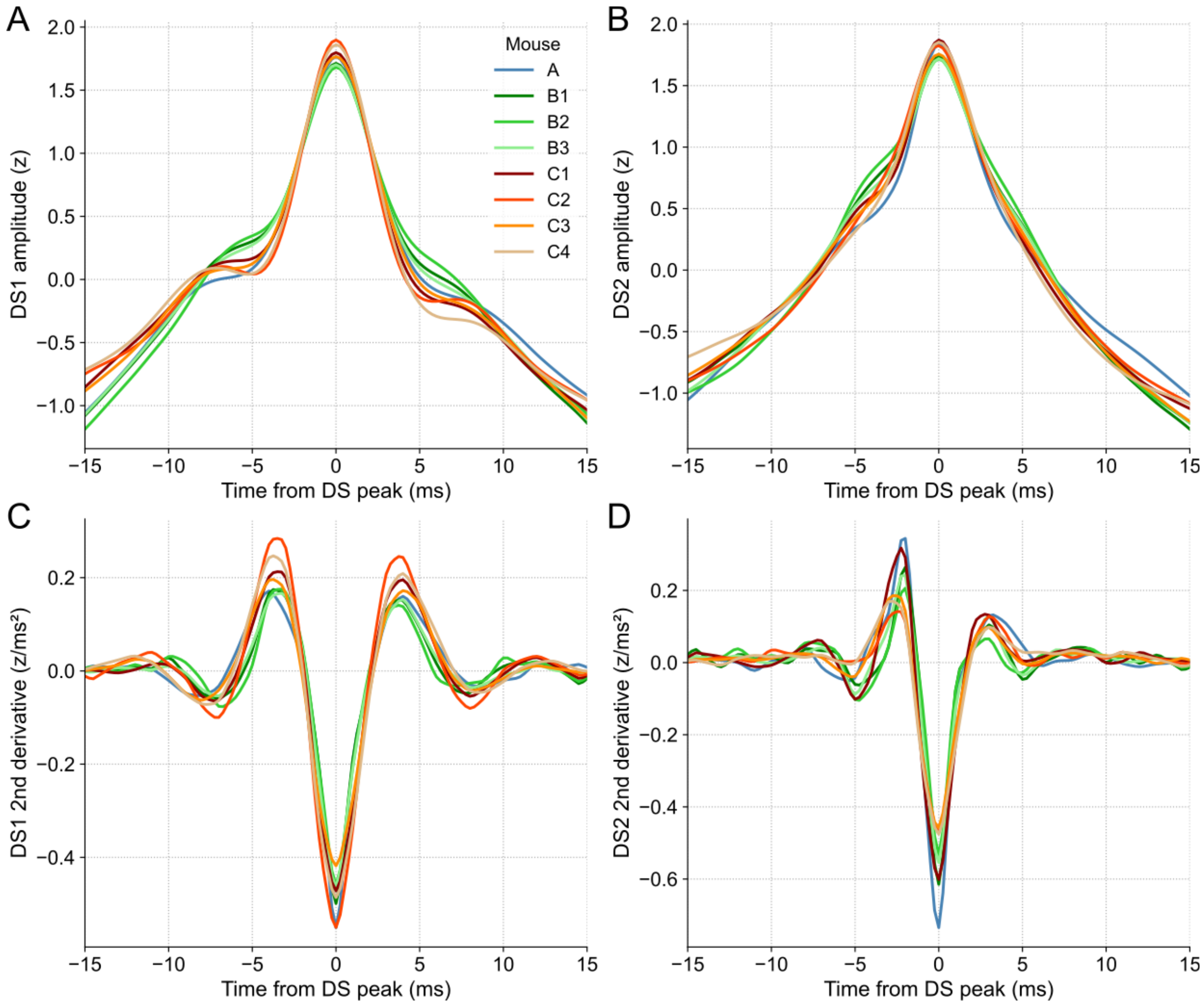


**Fig S4. Average waveform dynamics of each DS type of all mice.**

**(A)** Scaled mean waveforms of DS1 detected in the channel that yielded more DSs for all mice. **(B)** Same as A, but for DS2 **(C)** Second derivatives of the DS1 waveforms depicted in A. **(D)** Same as C, but for DS2. DS1: dentate spike type 1; DS2: dentate spike type 2.

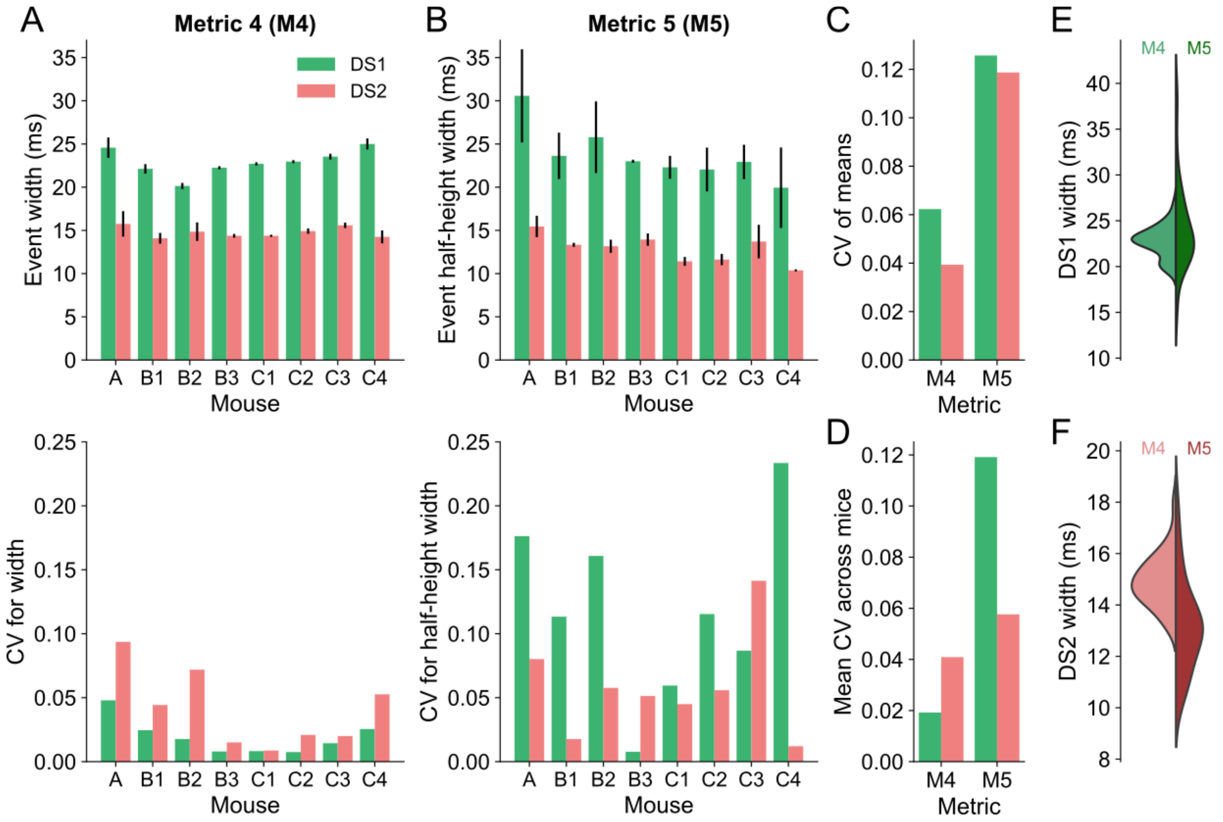


**Fig S5. DS width based on waveform dynamics is more stable than width at half height.**

**(A)** The top panel shows the width for each DS type (Metric 4 – M4) across channels for each mouse. Colored bars correspond to the means, and the error bars indicate the standard deviations. The bottom panel shows the coefficient of variation of the corresponding measurements in the panel above. **(B)** Same as A, but for the width at half-height (Metric 5 – M5). **(C)** Coefficients of variation of the means of each metric across mice. **(D)** Mean coefficients of variation of each metric across mice. **(E)** Distributions of M4 and M5 of DS1 showing that M5 is more variable. **(F)** Same as E, but for DS2. CV: coefficient of variation; DS: dentate spike; DS1: dentate spike type 1; DS2: dentate spike type 2.

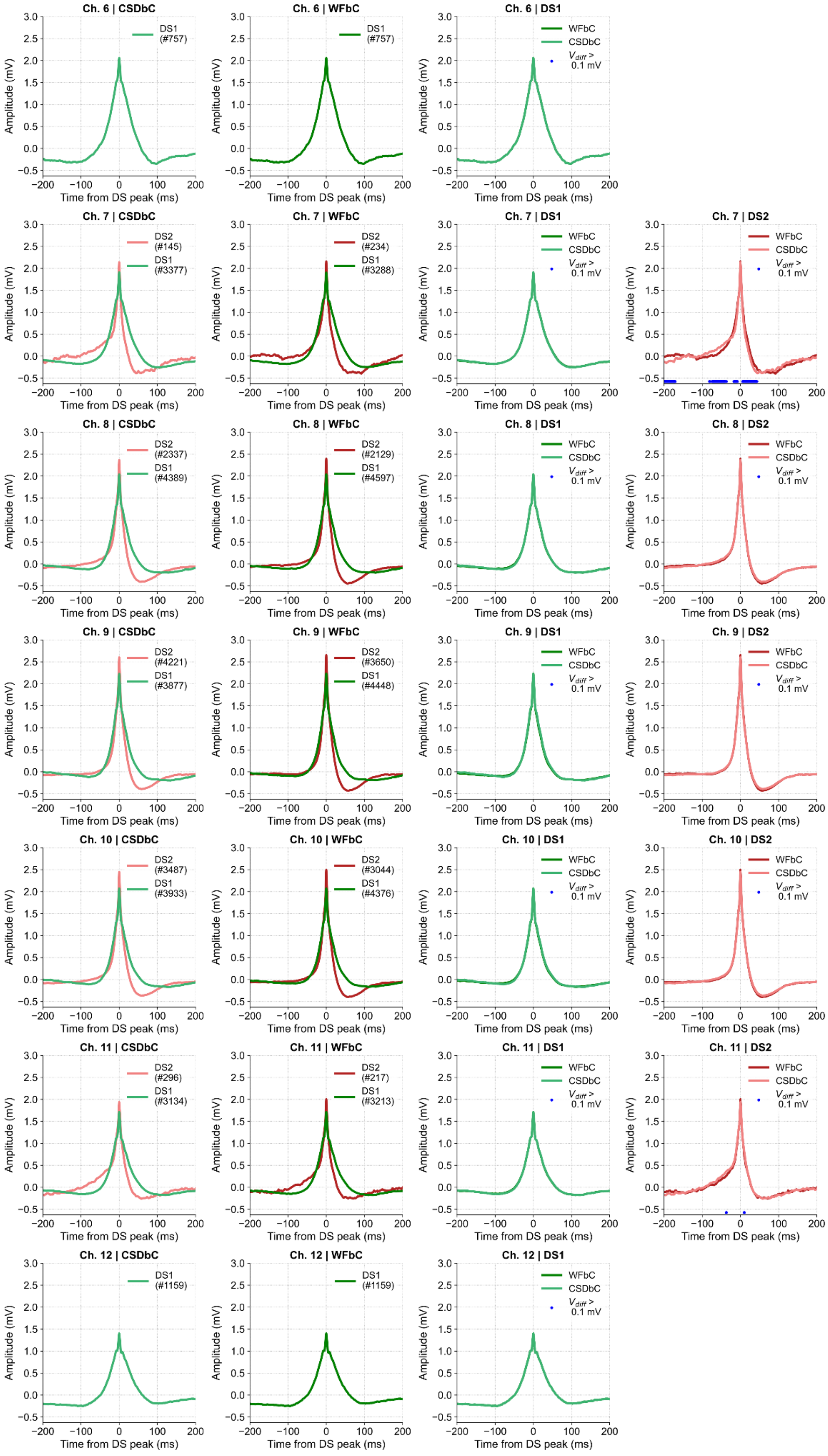


**Fig S6. DS waveform comparison between methods for each channel of mouse A.**

The panels in the first and second columns show the waveforms of each DS type classified by CSDbC and WFbC respectively. The panels in the third and fourth columns compare the waveforms of DS1 and DS2 respectively. Each line represents the chosen channel for DS detection. CSD: current source density; CSDbC: CSD-based classification; DS: dentate spike; DS1: DS type 1; DS2: DS type 2; WFbC: waveform-based classification.

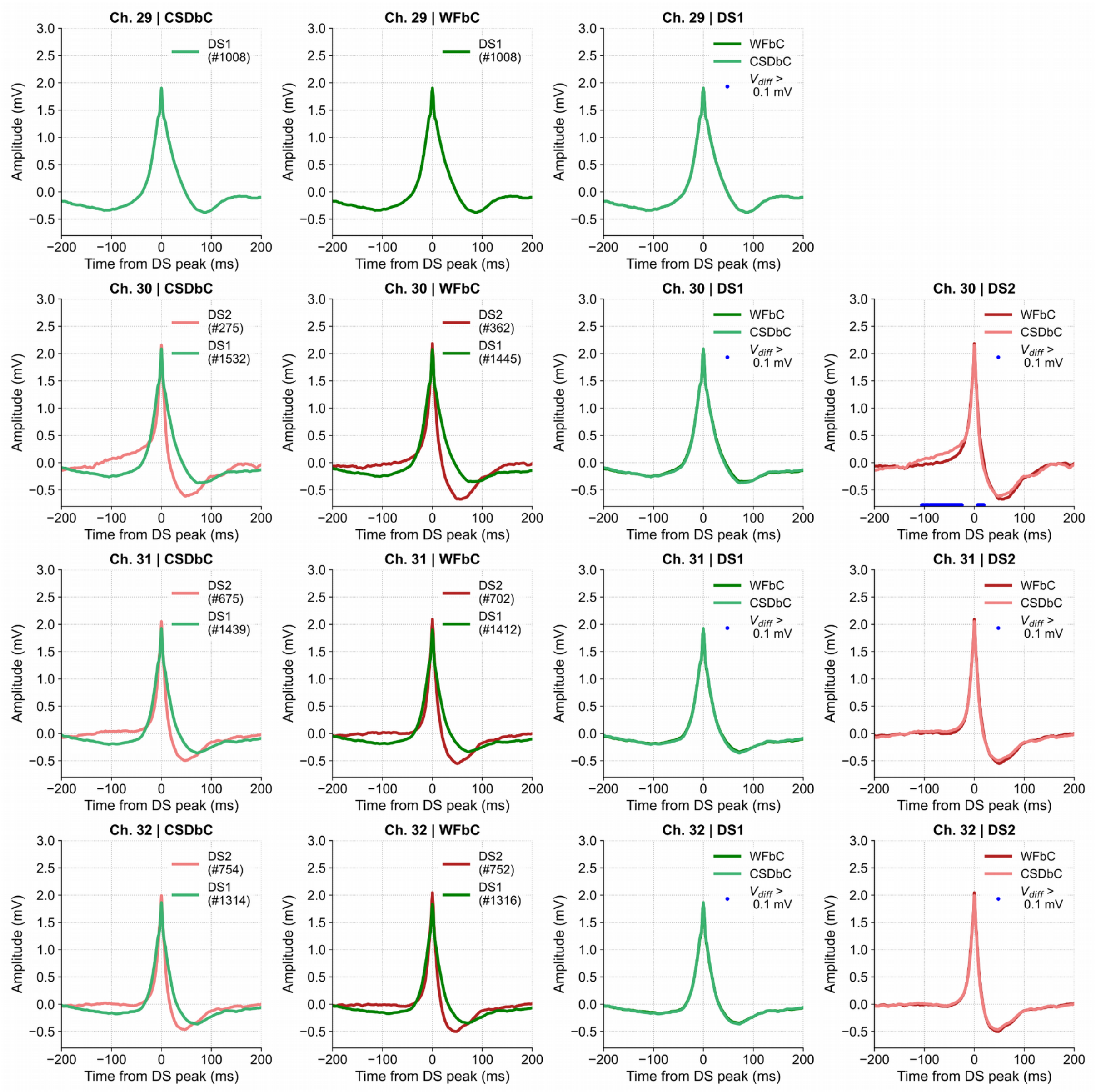


**Fig S7. DS waveform comparison between methods for each channel of mouse B1.**

The panels in the first and second columns show the waveforms of each DS type classified by CSDbC and WFbC respectively. The panels in the third and fourth columns compare the waveforms of DS1 and DS2 respectively. Each line represents the chosen channel for DS detection. CSD: current source density; CSDbC: CSD-based classification; DS: dentate spike; DS1: DS type 1; DS2: DS type 2; WFbC: waveform-based classification.

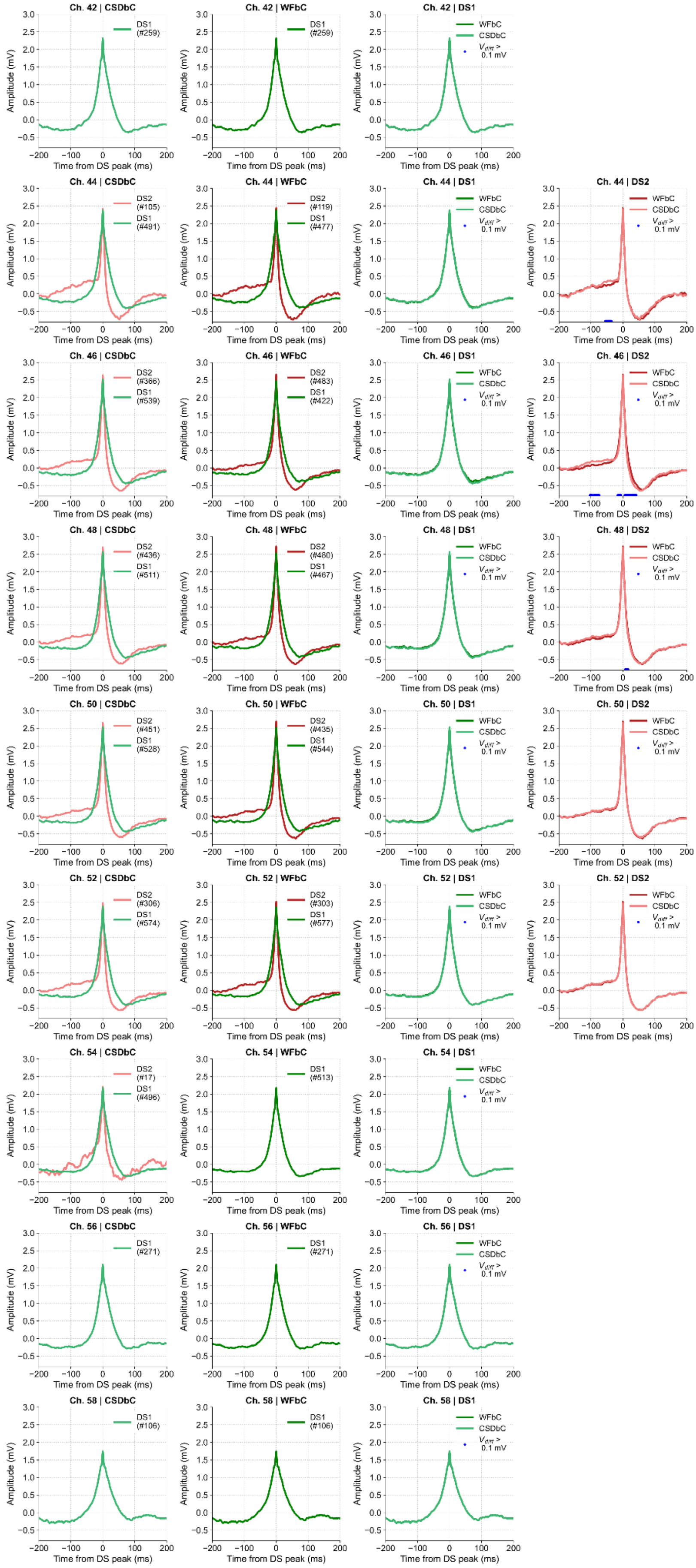


**Fig S8. DS waveform comparison between methods for each channel of mouse B2.** The panels in the first and second columns show the waveforms of each DS type classified by CSDbC and WFbC respectively. The panels in the third and fourth columns compare the waveforms of DS1 and DS2 respectively. Each line represents the chosen channel for DS detection. CSD: current source density; CSDbC: CSD-based classification; DS: dentate spike; DS1: DS type 1; DS2: DS type 2; WFbC: waveform-based classification.

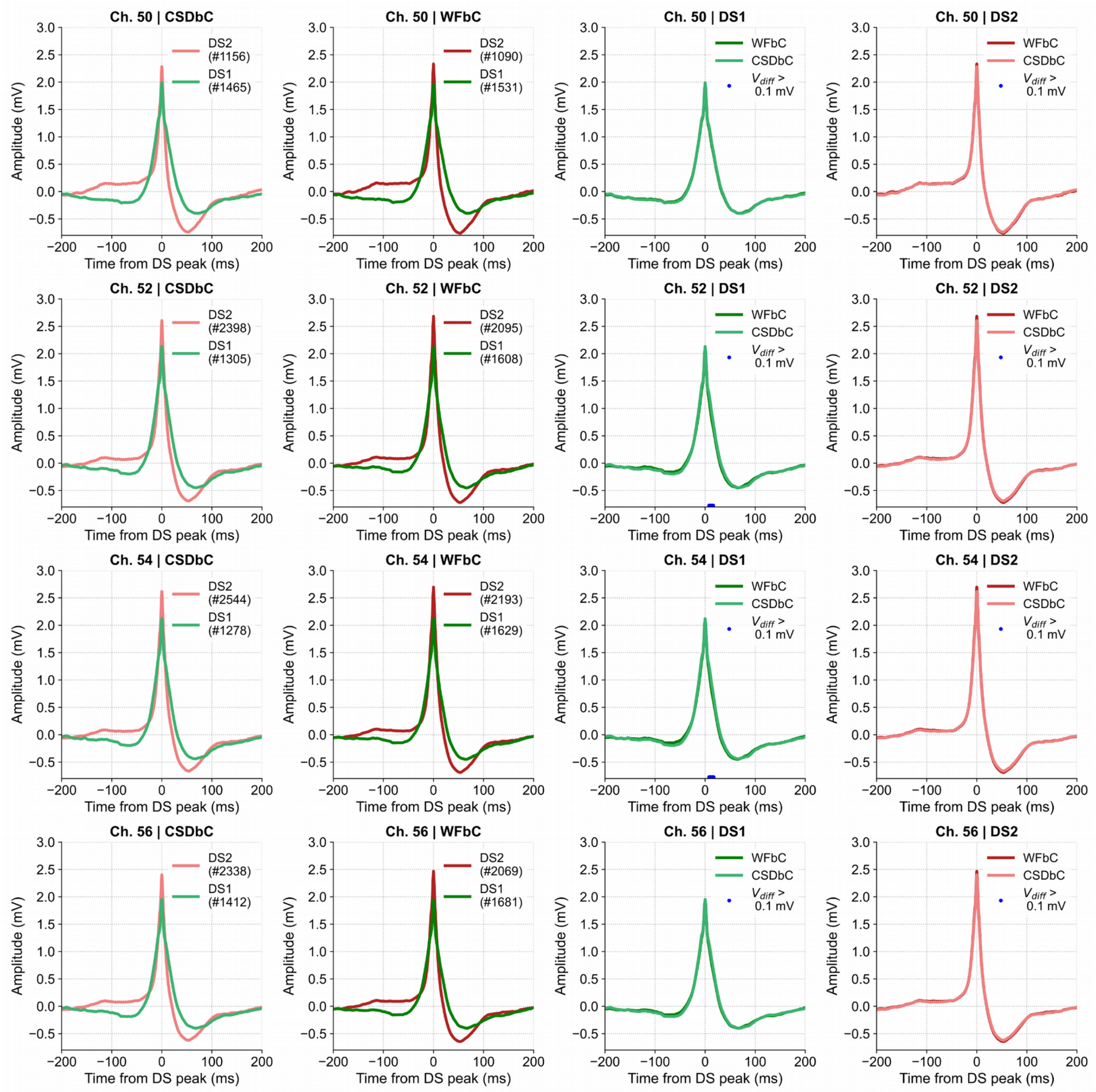


**Fig S9. DS waveform comparison between methods for each channel of mouse B3.**

The panels in the first and second columns show the waveforms of each DS type classified by CSDbC and WFbC respectively. The panels in the third and fourth columns compare the waveforms of DS1 and DS2 respectively. Each line represents the chosen channel for DS detection. CSD: current source density; CSDbC: CSD-based classification; DS: dentate spike; DS1: DS type 1; DS2: DS type 2; WFbC: waveform-based classification.

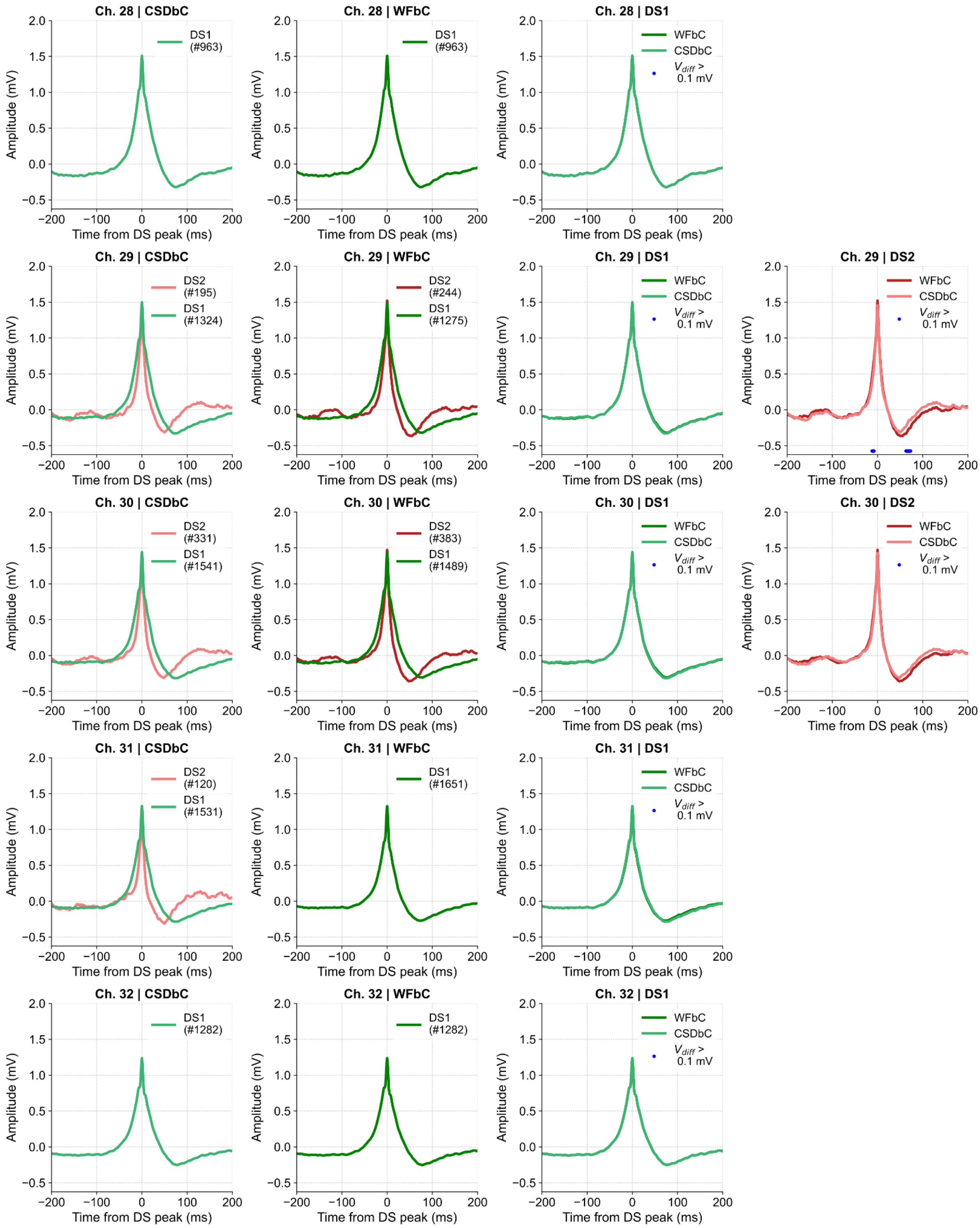


**Fig S10. DS waveform comparison between methods for each channel of mouse C1.**

The panels in the first and second columns show the waveforms of each DS type classified by CSDbC and WFbC respectively. The panels in the third and fourth columns compare the waveforms of DS1 and DS2 respectively. Each line represents the chosen channel for DS detection. CSD: current source density; CSDbC: CSD-based classification; DS: dentate spike; DS1: DS type 1; DS2: DS type 2; WFbC: waveform-based classification.

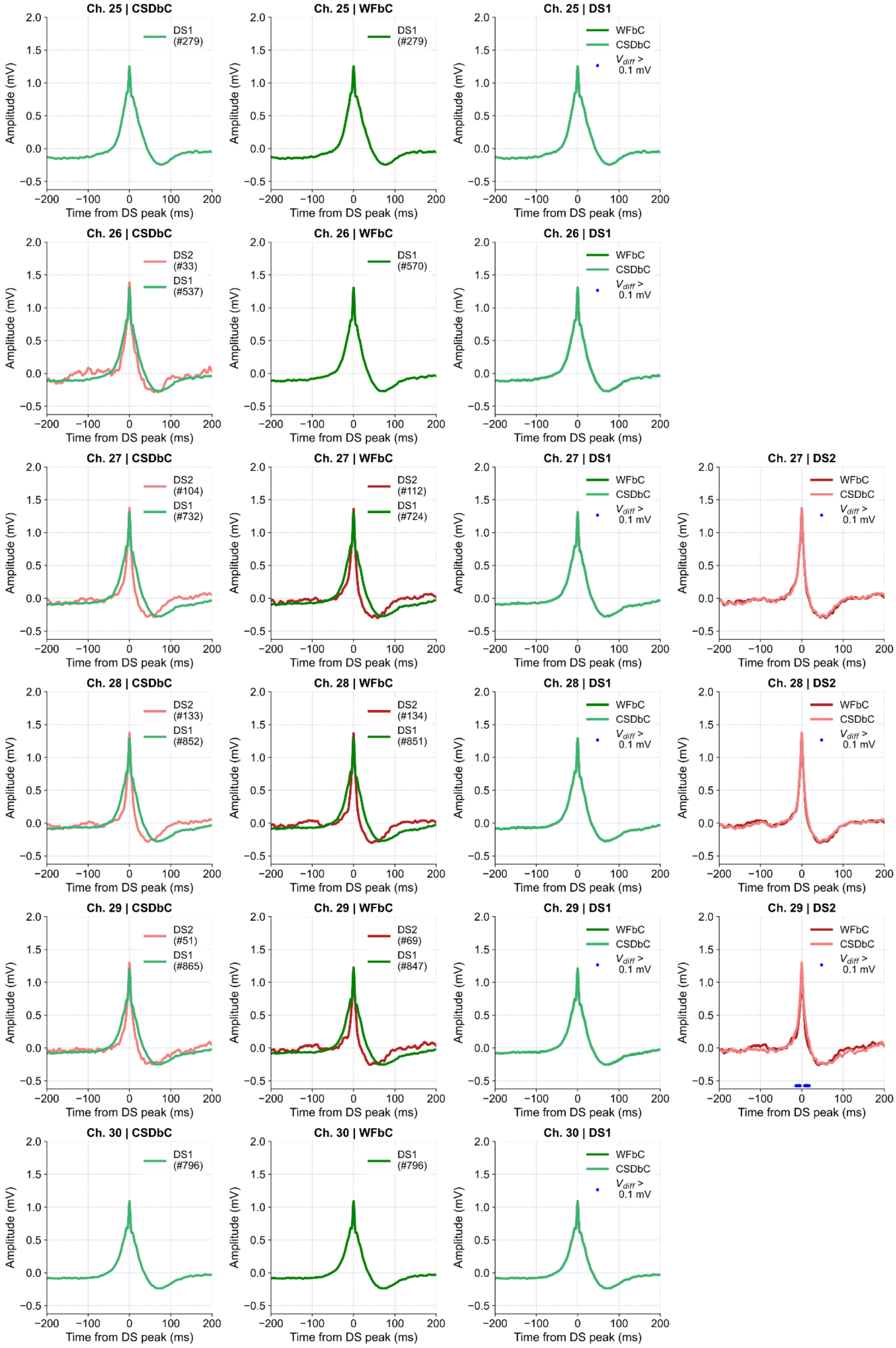


**Fig S11. DS waveform comparison between methods for each channel of mouse C2.**

The panels in the first and second columns show the waveforms of each DS type classified by CSDbC and

WFbC respectively. The panels in the third and fourth columns compare the waveforms of DS1 and DS2 respectively. Each line represents the chosen channel for DS detection. CSD: current source density; CSDbC: CSD-based classification; DS: dentate spike; DS1: DS type 1; DS2: DS type 2; WFbC: waveform-based classification.

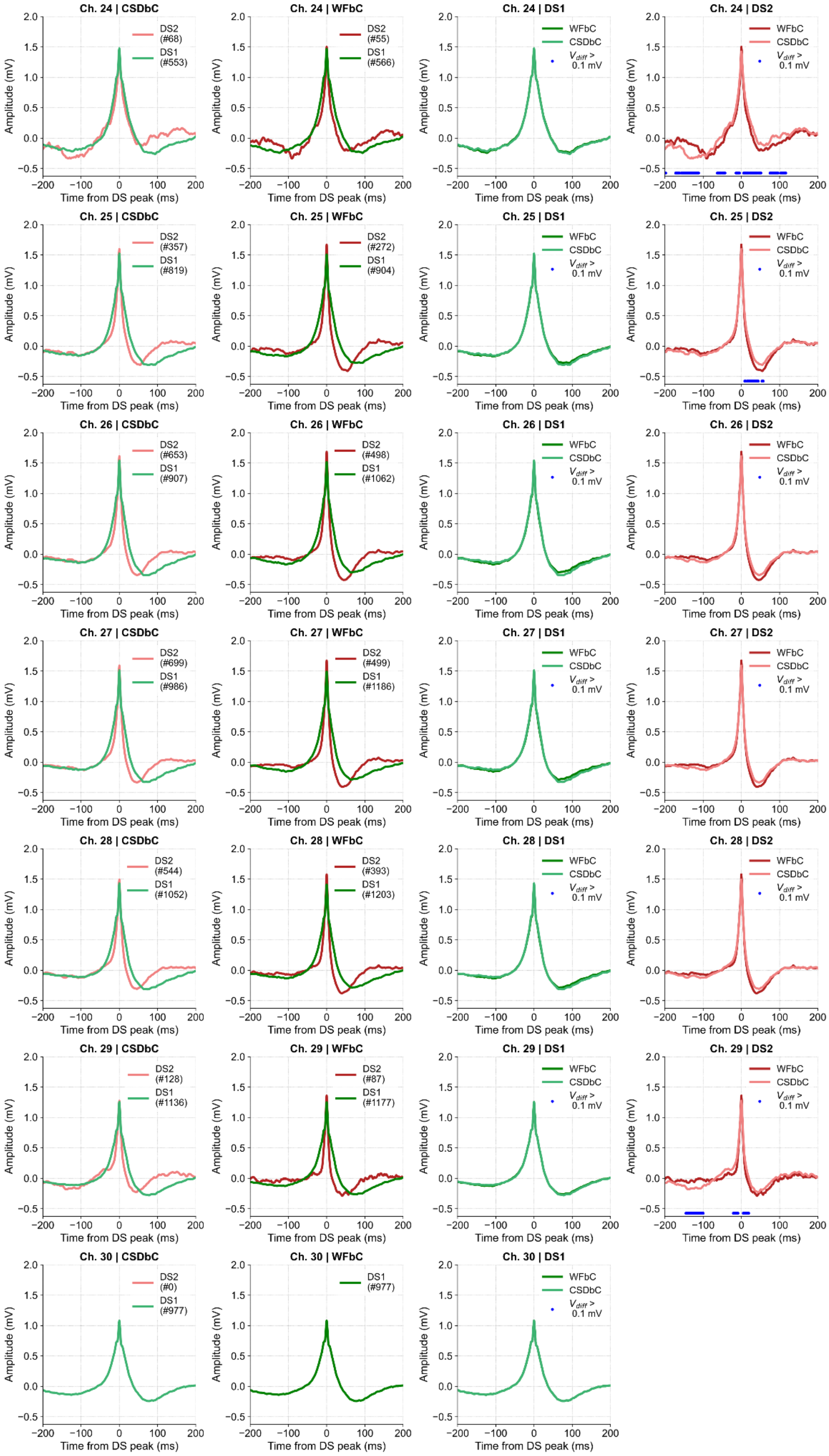

**Fig S12. DS waveform comparison between methods for each channel of mouse C3.**

The panels in the first and second columns show the waveforms of each DS type classified by CSDbC and WFbC respectively. The panels in the third and fourth columns compare the waveforms of DS1 and DS2 respectively. Each line represents the chosen channel for DS detection. CSD: current source density; CSDbC:

CSD-based classification; DS: dentate spike; DS1: DS type 1; DS2: DS type 2; WFbC: waveform-based classification.

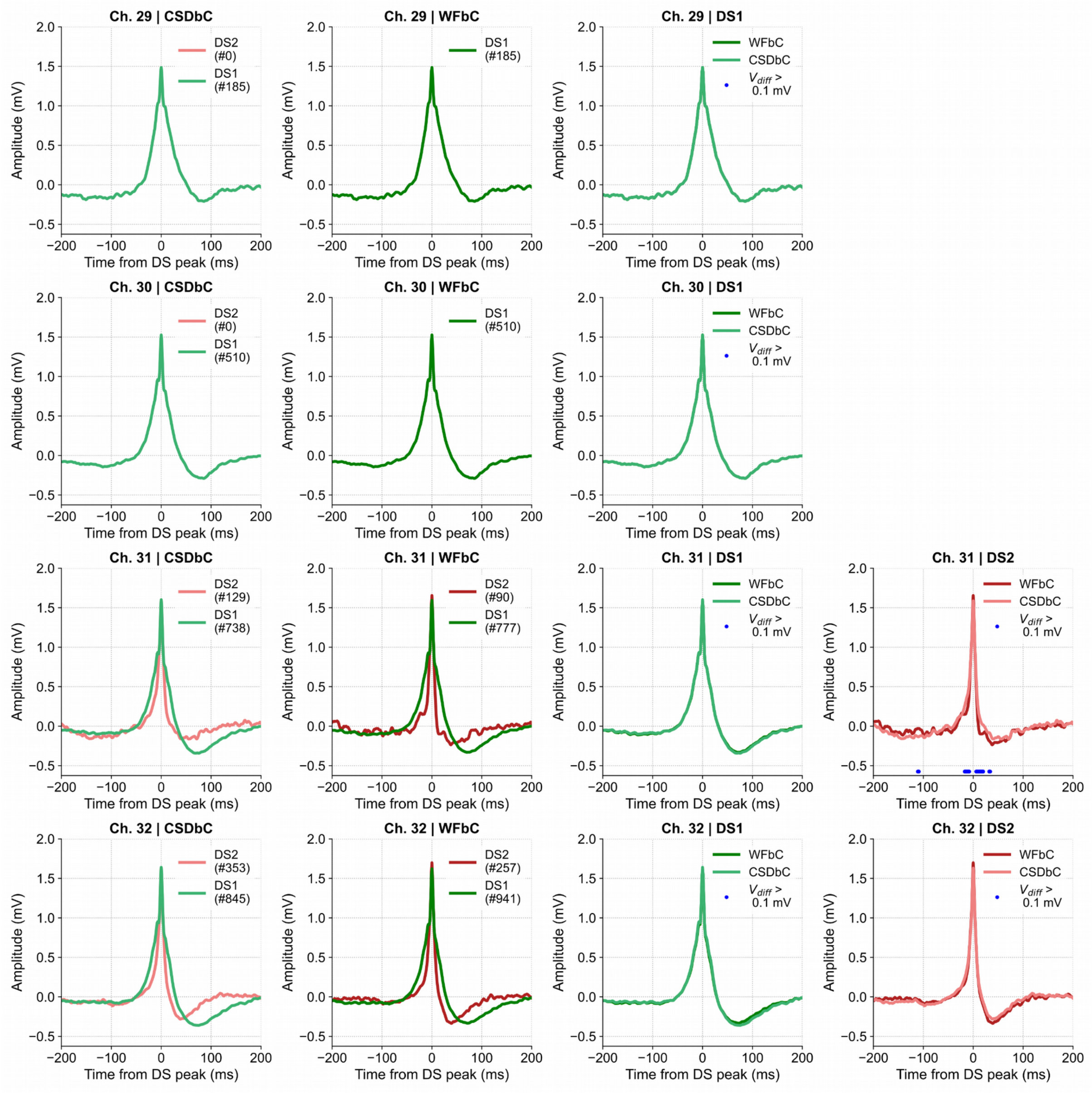


**Fig S13. DS waveform comparison between methods for each channel of mouse C4.**

The panels in the first and second columns show the waveforms of each DS type classified by CSDbC and WFbC respectively. The panels in the third and fourth columns compare the waveforms of DS1 and DS2 respectively. Each line represents the chosen channel for DS detection. CSD: current source density; CSDbC: CSD-based classification; DS: dentate spike; DS1: DS type 1; DS2: DS type 2; WFbC: waveform-based classification.

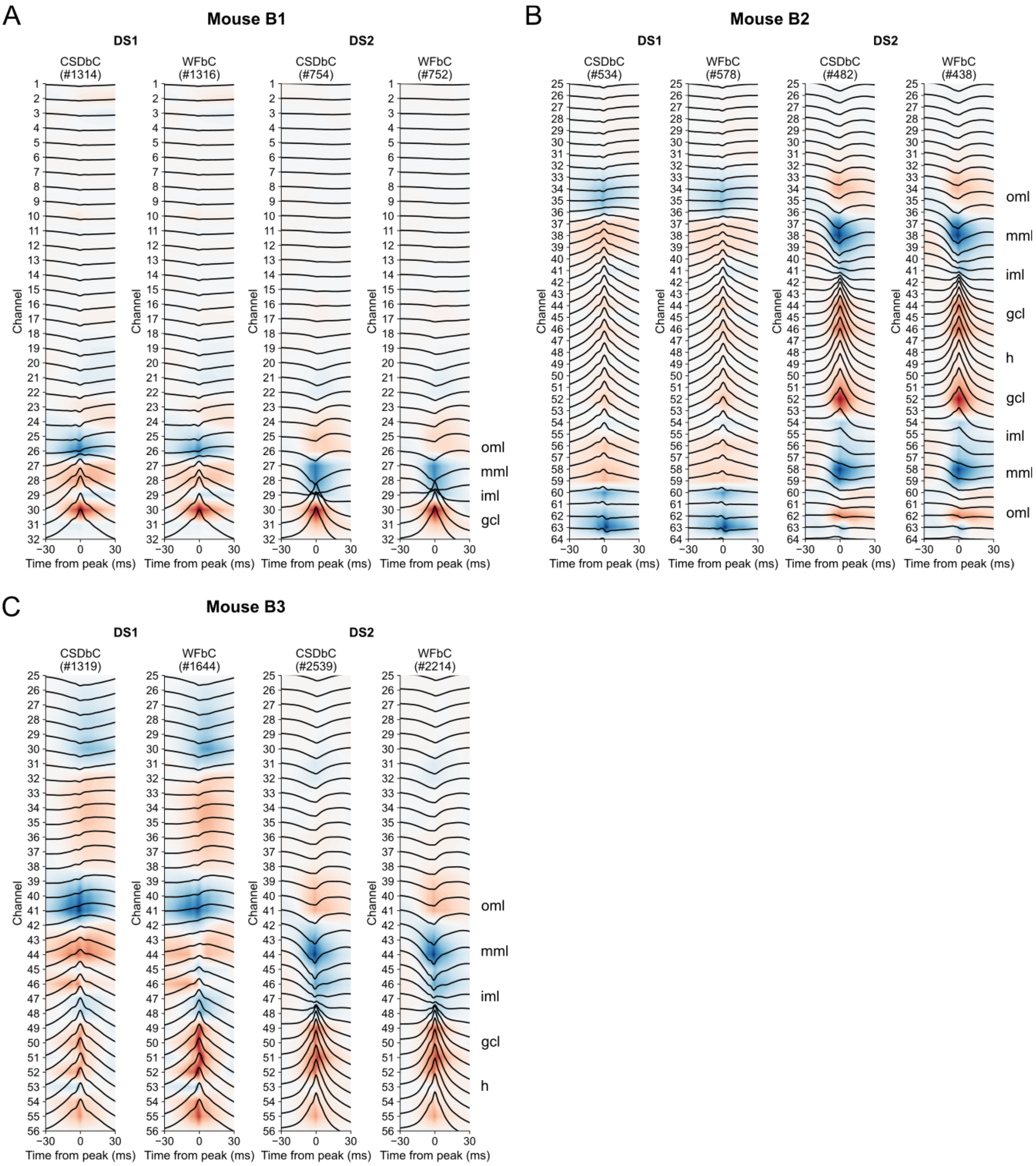


**Fig S14. DS-triggered CSDs from mice of dataset B.**

**(A)** Mean CSDs of mouse B1 within a 60-ms window around the peak of each DS type classified by CSDbC and WFbC. Blue and red colors represent sinks and sources, respectively. The estimated DG layers are indicated on the right. **(B,C)** Same as A, but for mice B2 and B3. CSD: current source density; CSDbC: CSD-based classification; DS: dentate spike; DS1: DS type 1; DS2: DS type 2; WFbC: waveform-based classification; gcl: granule cell layer; h: hilus; iml: inner molecular layer; mml: middle molecular layer; oml: outer molecular layer.

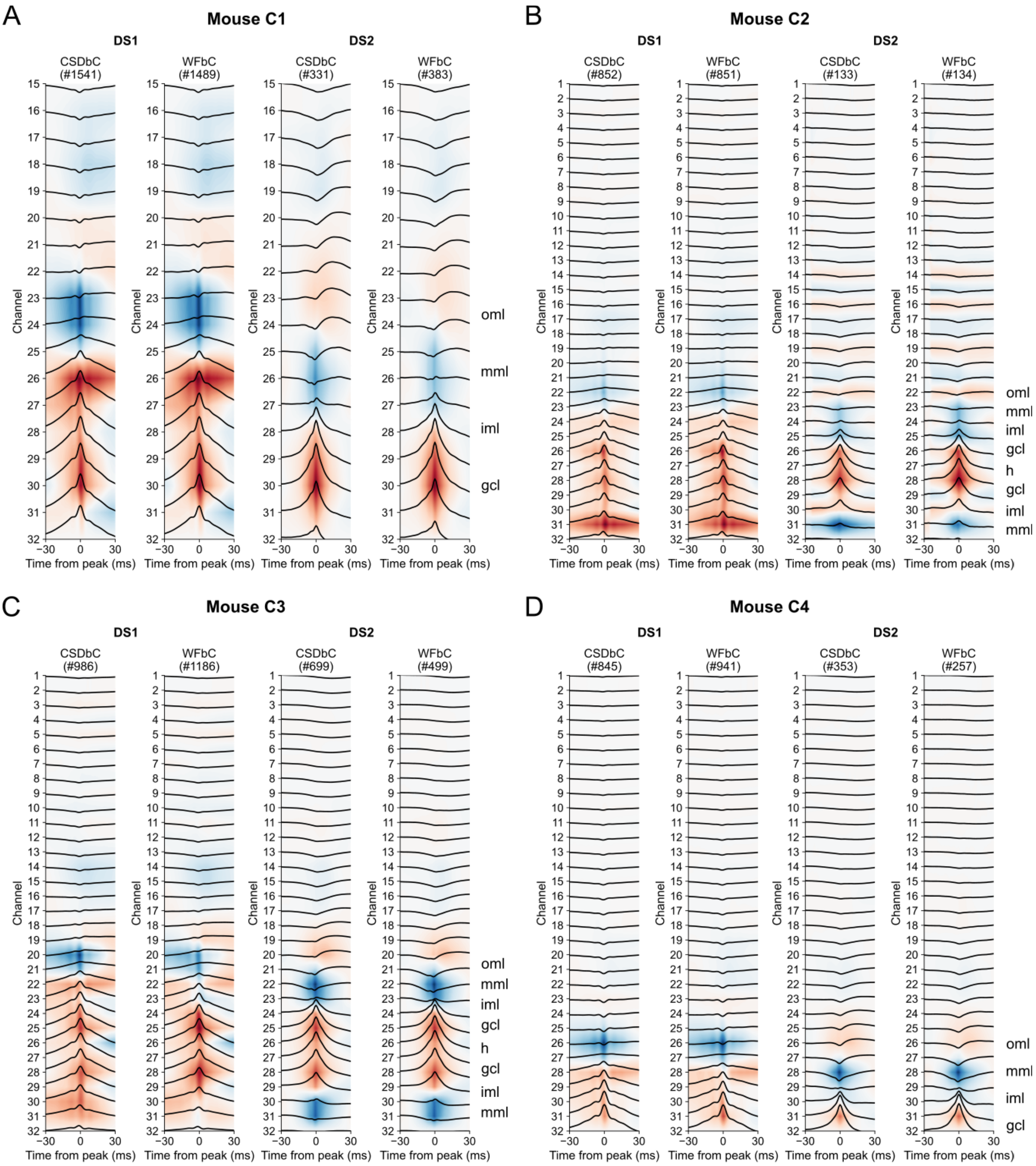


**Fig S15. DS-triggered CSDs from mice of dataset C.**

**(A)** Mean CSDs of mouse C1 within a 60-ms window around the peak of each DS type classified by CSDbC and WFbC. Blue and red colors represent sinks and sources, respectively. The estimated DG layers are indicated on the right. **(B-D)** Same as A, but for mice C2, C3 and C4. CSD: current source density; CSDbC: CSD-based classification; DS: dentate spike; DS1: DS type 1; DS2: DS type 2; WFbC: waveform-based classification; gcl: granule cell layer; h: hilus; iml: inner molecular layer; mml: middle molecular layer; oml: outer molecular layer.

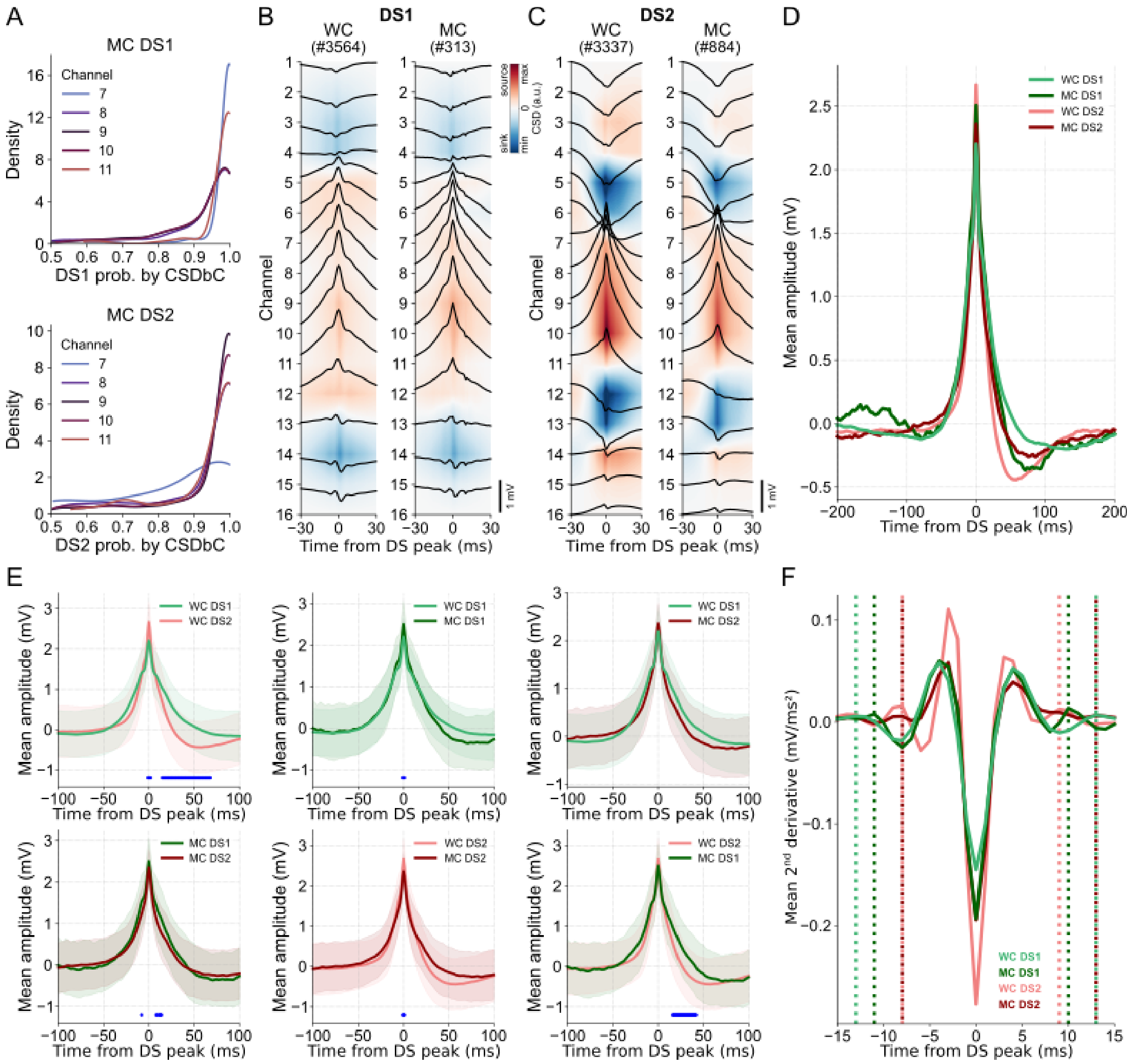


**Fig S16. Misclassified DSs exhibit post-peak dynamics similar to related types.**

**(A)** The probability densities of DS1 misclassified as DS2 by WFbC in channels 7 to 11 of mouse A is shown on top. The bottom panel shows the same for DS2. **(B)** Mean CSDs for the well-classified (left panel) and misclassified (right panel) DS1 by WFbC. **(C)** Same as B, but for DS2. **(D)** Mean waveforms of well- and misclassified DSs of both types in channel 9. **(E)** Pair-to-pair comparison of the waveforms shown in D. The blue dots indicate when Cohen's D size effect is greater than 0.5 for periods with significant differences ($p$ < 0.05 in t-test) between the waveforms. **(F)** Mean 2$^{nd}$ derivatives of the waveforms shown in D. The dotted lines indicate the start and end width limits of each waveform. DS1: dentate spike type 1; DS2: dentate spike type 2; MC: misclassified; WC: well-classified.

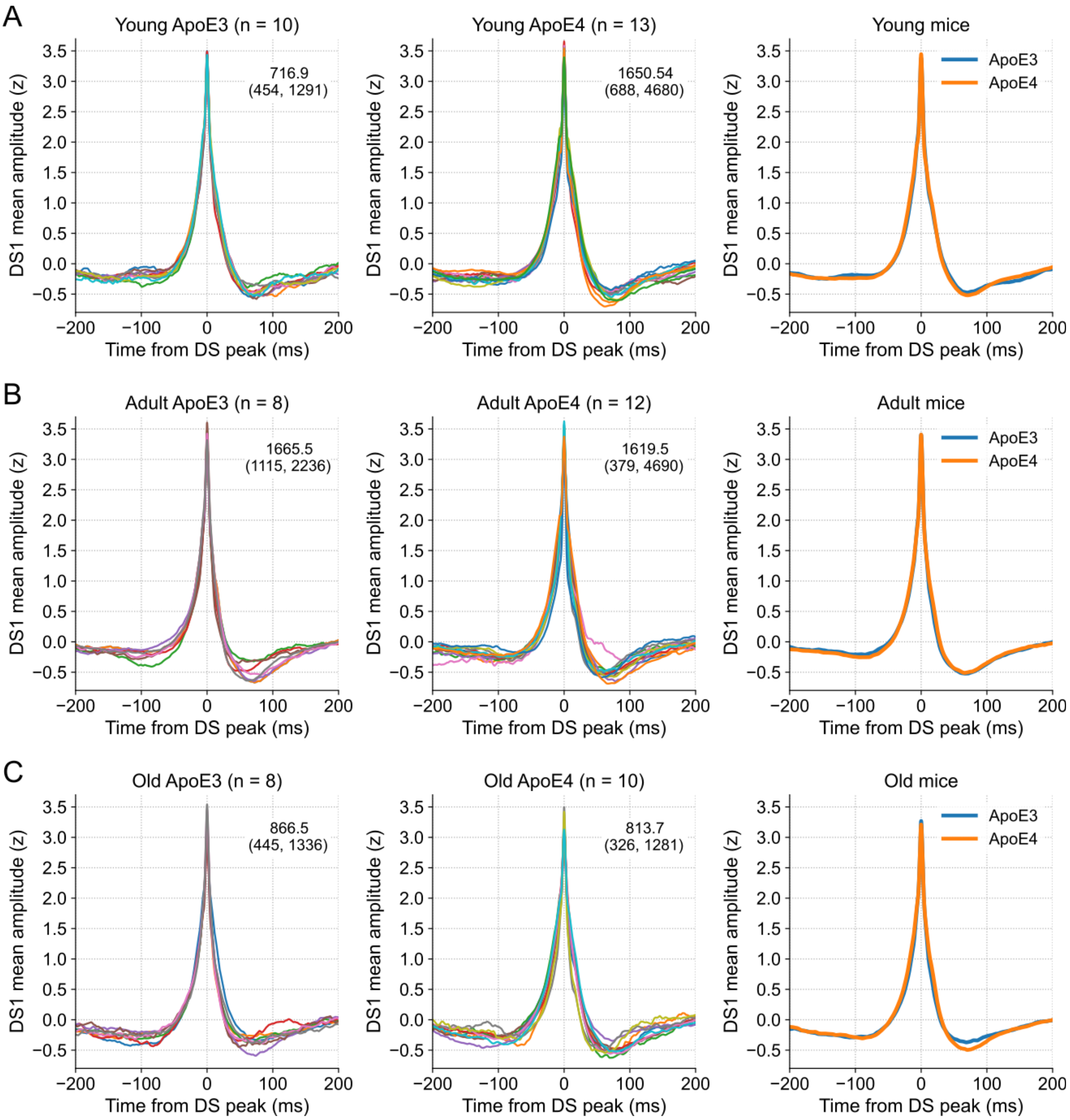


**Fig S17. DS1 mean waveforms of ApoE3-KI and ApoE4-KI mice.**

**(A)** DS1 mean waveforms of each ApoE3-KI mouse (left panel) and each ApoE4-KI mouse (middle panel) when young. Numbers within the graphs indicate the average number of events with minimum and maximum values in parentheses. The comparison of overall mean waveforms is shown in the right panel. **(B,C)** Same as A, but for mice in adult and old ages. ApoE3-KI: Apolipoprotein E3 knock-in mice; ApoE4-KI: Apolipoprotein E4 knock-in mice; DS1: dentate spike type 1; DS2: dentate spike type 2.

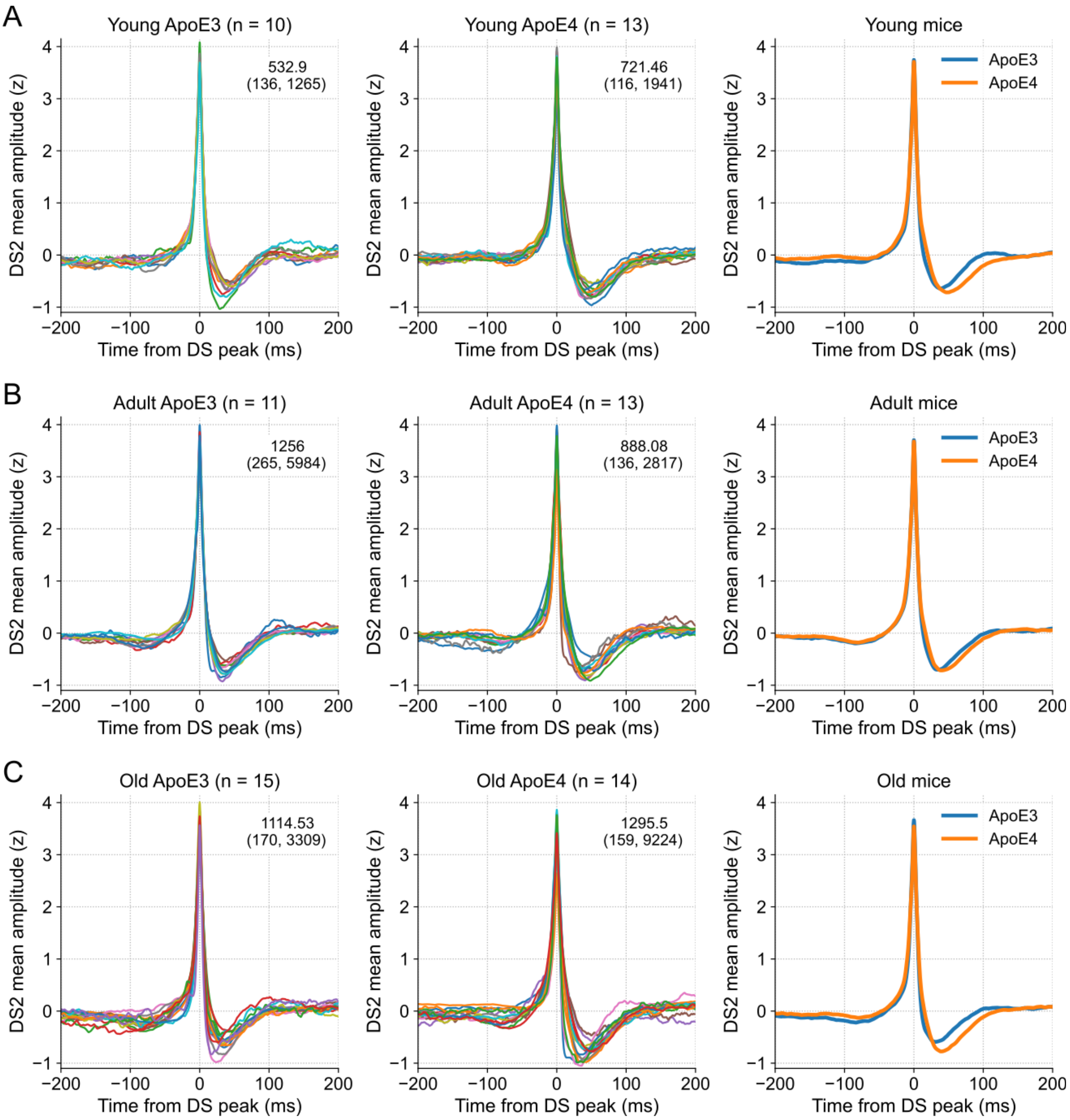


**Fig S18. DS2 mean waveforms of ApoE3-KI and ApoE4-KI mice.**

**(A)** DS2 mean waveforms of each ApoE3-KI mouse (left panel) and each ApoE4-KI mouse (middle panel) when young. Numbers within the graphs indicate the average number of events with minimum and maximum values in parentheses. The comparison of overall mean waveforms is shown in the right panel. **(B,C)** Same as A, but for mice in adult and old ages. ApoE3-KI: Apolipoprotein E3 knock-in mice; ApoE4-KI: Apolipoprotein E4 knock-in mice; DS1: dentate spike type 1; DS2: dentate spike type 2.

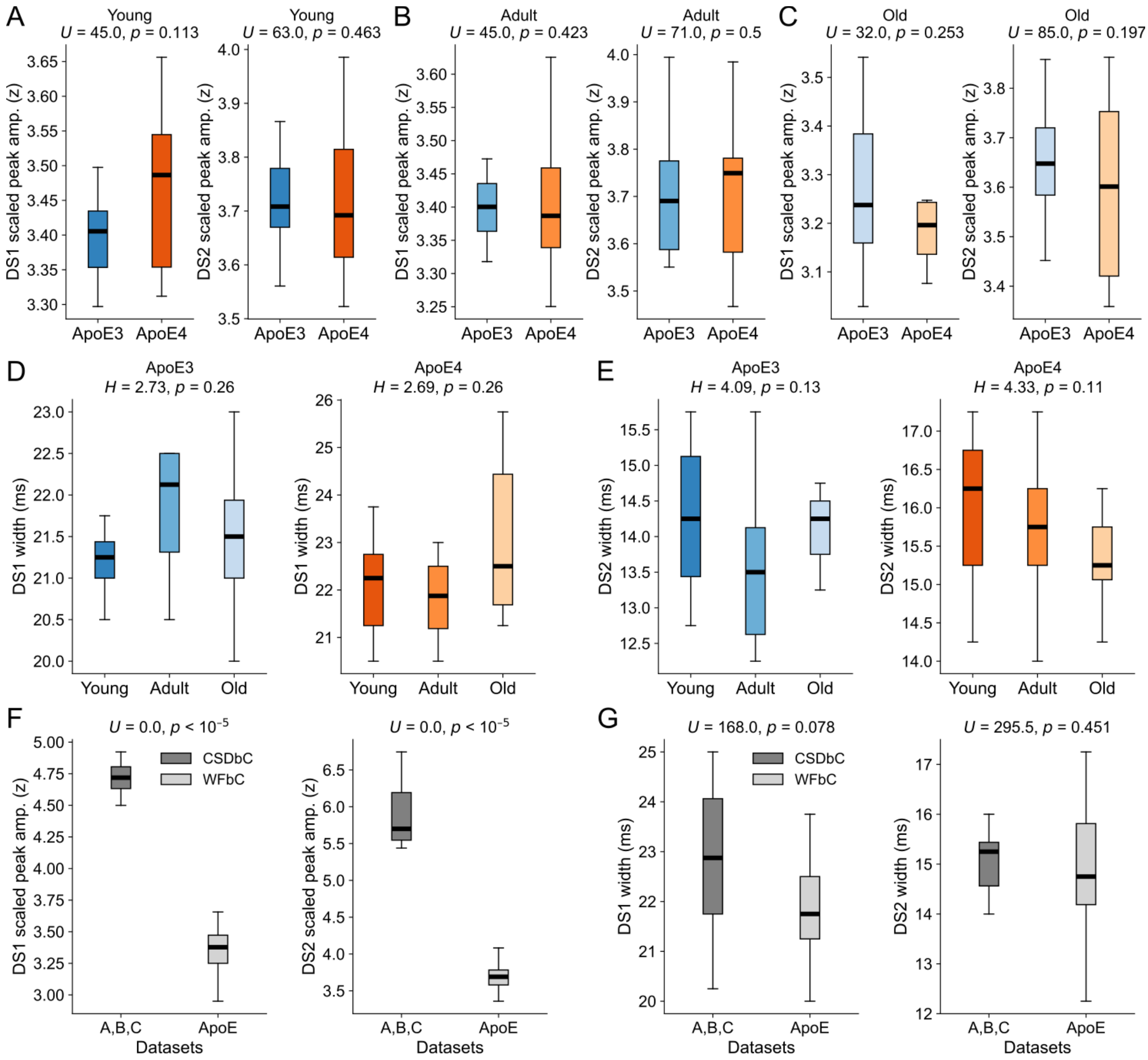


**Fig S19. Stable DS metrics across genotypes, ages and datasets.**

**(A)** Comparison of the scaled peak amplitudes of DS1 (left panel) and DS2 (right panel) between young ApoE3-KI and ApoE4-KI mice. **(B,C)** Same as A, but for mice in adult and old ages. **(D)** Comparison of DS1 widths of ApoE3-KI (left panel) and ApoE4-KI mice (right panel) across ages. **(E)** Same as D, but for DS2. **(F)** Comparison of scaled peak amplitudes of DS1 (left panel) and DS2 (right panel) between mice from datasets A-C and ApoE transgenic mice, whose DSs were classified via CSDbC and WFbC respectively. **(G)** Same as F, but for DS widths. Statistics and *p*-values were obtained from Mann-Whitney *U* tests for panels in A-C,F,G, and from Kruskal-Wallis H tests for panels in D and E. ApoE3-KI: Apolipoprotein E3 knock-in mice; ApoE4-KI: Apolipoprotein E4 knock-in mice; DS1: dentate spike type 1; DS2: dentate spike type 2.
